# Mutual repression between JNK/AP-1 and JAK/STAT stratifies cell behaviors during tissue regeneration

**DOI:** 10.1101/2022.05.11.491445

**Authors:** Janhvi Jaiswal, Raphael Engesser, Andrea Armengol Peyroton, Vanessa Weichselberger, Carlo Crucianelli, Isabelle Grass, Jens Timmer, Anne-Kathrin Classen

## Abstract

Epithelial repair relies on the activation of stress signaling pathways to coordinate cellular repair behaviors. Their deregulation is implicated in chronic wound and cancer pathologies. Despite such translational importance, an understanding of how spatial patterns of signaling pathways and repair behaviors arise in damaged tissues remains elusive. Using TNF-α/Eiger-mediated inflammatory damage to *Drosophila* imaginal discs, we uncover that JNK/AP-1 signaling cells act as paracrine organizers and initiate a mutual repression network that spatially segregates JNK/AP-1 and JAK/STAT signaling cells into distinct populations. While JNK/AP-1 signaling cells produce JAK/STAT-activating Upd ligands, these signal-sending cells suppress activation of JAK/STAT via Ptp61F. Conversely, responding cells with activated JAK/STAT suppress JNK activation via Zfh2. The resulting bistable segregation of signaling domains is associated with distinct cellular tasks and regenerative potential. While JNK/AP-1 signaling cells at the wound center act as paracrine organizers, their cell cycle is senescently arrested. Thus, compensatory proliferation occurs exclusively in JAK/STAT signaling cells at the wound periphery. This spatial stratification is essential for proper tissue repair, as co-activation of JNK/AP-1 and JAK/STAT in the same cells creates conflicting inputs on cell cycle progression, leading to excess apoptosis of senescently arrested organizer cells. Finally, we demonstrate that bistable spatial segregation of JNK/AP-1 and JAK/STAT drives senescent and proliferative behaviors in transient as well as chronic tissue damage models, and importantly, in *Ras^V12^, scrib* tumors under the influence of JNK/AP-1 activity. Revealing this previously uncharacterized regulatory network between JNK/AP-1, JAK/STAT and associated cell behaviors have important implications for our conceptual understanding of tissue repair, chronic wound pathologies and tumor microenvironments, where both pathways are strongly implicated.

## Introduction

Wound repair programs rely on the coordination of distinct responses to damage [1, 2]. Upon damage, wound-derived factors initiate inflammation and stress-signaling pathways which drive cellular responses like apoptosis, proliferation, survival and tissue remodeling [3–6]. While these responses are essential, it is critical that they remain spatio-temporally restricted to avoid the establishment of chronic wounds [7–9]. Chronic wounds are characterized by sustained inflammation, and deregulated proliferation and apoptosis [10]. The finding that these are also hallmarks of tumor microenvironments supported the idea that tumors resemble chronic, non-healing wounds [11–13]. Yet, the tissue-level coordination of signaling pathways and repair behaviors upon wounding, and the conditions under which they turn pathological remain elusive.

A myriad of wound models identified specific signaling pathways required for regeneration, of which several have also been shown to drive tumor growth. In particular, the JNK/AP-1 and the JAK/STAT pathways are consistently implicated in regeneration [14–17] as well as in tumors [18–23]. In the context of wound-healing, JNK/AP-1 activation is indispensable for regeneration [24–26]. JNK/AP-1 is one of the earliest signaling responses to damage, proposedly activated by wound-derived ROS signals [27, 28]. In this role, JNK/AP-1 regulates a variety of conflicting cell behaviors including apoptosis [29, 30], survival [31] and compensatory proliferation [32, 33]. These paradoxical behaviors have been extensively characterized individually [34–36], yet how they are organized on a tissue scale to ensure regeneration is not known.

We recently reported that high JNK/AP-1 signaling facilitates survival in wounds and tumors by mediating a cell cycle stall in G2 which is characterized by anti-apoptotic and senescent features [31]. Forced exit of these cells from G2 into mitosis and G1 caused a substantial increase in apoptosis, suggestive of the JNK-mediated G2 stall being a protective state during regeneration (Fig. S1A) [31]. This model is supported by recent evidence that the universal integrator of molecular damage, p53, is activated by the G2/M kinase Cdk1 to induce apoptosis competence upon exit from G2 [37]. The G2 protective state is critical as high JNK/AP-1 signaling cells are shown to produce mitogens necessary to induce compensatory proliferation and regeneration [28, 38–40]. Paradoxically, this reveals a cell population at the center of wounds which produces pro-proliferative signals but itself is prohibited from proliferation. These findings highlight the necessity of a mechanism, which ensures that high JNK/AP-1 signaling cells maintain a G2 stall in the presence of mitogenic signals, thereby outsourcing proliferation to other cells. Alternatively, it requires the existence of a mechanism that promotes cycling as well as survival of JNK-signaling cells. This illustrates that the underlying mechanisms and molecular effectors that resolve these conflicting cell behaviors downstream of JNK/AP-1 signaling in the same tissues remains elusive.

Prominent cytokines produced by JNK/AP-1 signaling cells belong to the Unpaired (Upd) family [41–45] and activate the JAK/STAT pathway which, in turn, plays a significant role in mediating proliferation [18, 20, 28, 39, 46–50] and cell survival during tissue stress [51, 52]. Interestingly, we have previously also shown that JAK/STAT is required in the disc to restrict the expansion of JNK/AP-1 signaling [51]. This is essential as JNK/AP-1 signaling within the tissue can self-propagate by promoting an increase in the expression of its own paracrine activators - the TNF-α/Eiger (Egr) cytokine [53, 54], as well as ROS [28, 55, 56].

These observations indicate an interdependence between the JNK/AP-1 and JAK/STAT signaling pathways, whose coexistence has been consistently reported in wounds [14–17] and tumor tissues [18–23]. Yet the regulatory network of the two pathways, beyond the regulation of *upd’s* by JNK, remains so far unexplored. As JNK/AP-1 and JAK/STAT drive contradicting yet critical cell behaviors - namely G2 stalling and proliferation, linked by their regulation of survival - characterizing their regulatory interactions is essential to understand how these pathways cooperatively organize tissue stress responses.

## Results

### Figure 1 JNK/AP-1 and JAK/STAT segregate into distinct spatial domains upon tissue damage

To characterize the JNK/AP-1 and JAK/STAT signaling pathways at wound sites in detail, we analyzed signaling dynamics and repair behaviors induced by targeted expression of the TNF-α homologue *eiger* (*egr*) in the wing disc pouch (Fig. S1.1B,C). Ectopic *egr*-expression has been extensively used as a model to genetically induce wounds and study regeneration [57]. *egr*-expression activates the stress-signaling pathway JNK/AP-1, assessed by the sensitive JNK/AP-1 reporter - *TRE>RFP* (Fig. 1A,B) [58], which leads to loss of polarity, epithelial barrier dysfunction (Fig. S1.1D-E’) and inflammation [24, 25, 59]. Specifically, expression of *egr* also induces *upd1-3* cytokine expression [51], MMP-1 production [60–62], ROS production [8, 63] and protective responses including UPR [64] and, ultimately, NF-κB signaling upregulation [65] (Fig.S1.1F-O, see also Fig.1E,F) in high JNK/AP-1 signaling cells – which are all known hallmarks of inflammatory, and by extension, chronic wounds [6, 10]. In addition, high JNK/AP-1 activity at the center of the *egr-*expressing domain (Fig. 1A,B) induces a senescent-like G2 cell cycle stall (Fig. 1C,D) which is essential for cell survival (Fig. S1.1A), as assessed using the Fly-FUCCI toolkit [31, 66]. Indeed, we found that proliferation was excluded from the inflammatory JNK-signaling domain at the center of the disc and was restricted to the wound periphery (Fig. 1C,D,N). These observations highlight a pronounced spatial segregation of regenerative cell behaviors linked to JNK-induced tissue damage responses.

**Figure 1.**
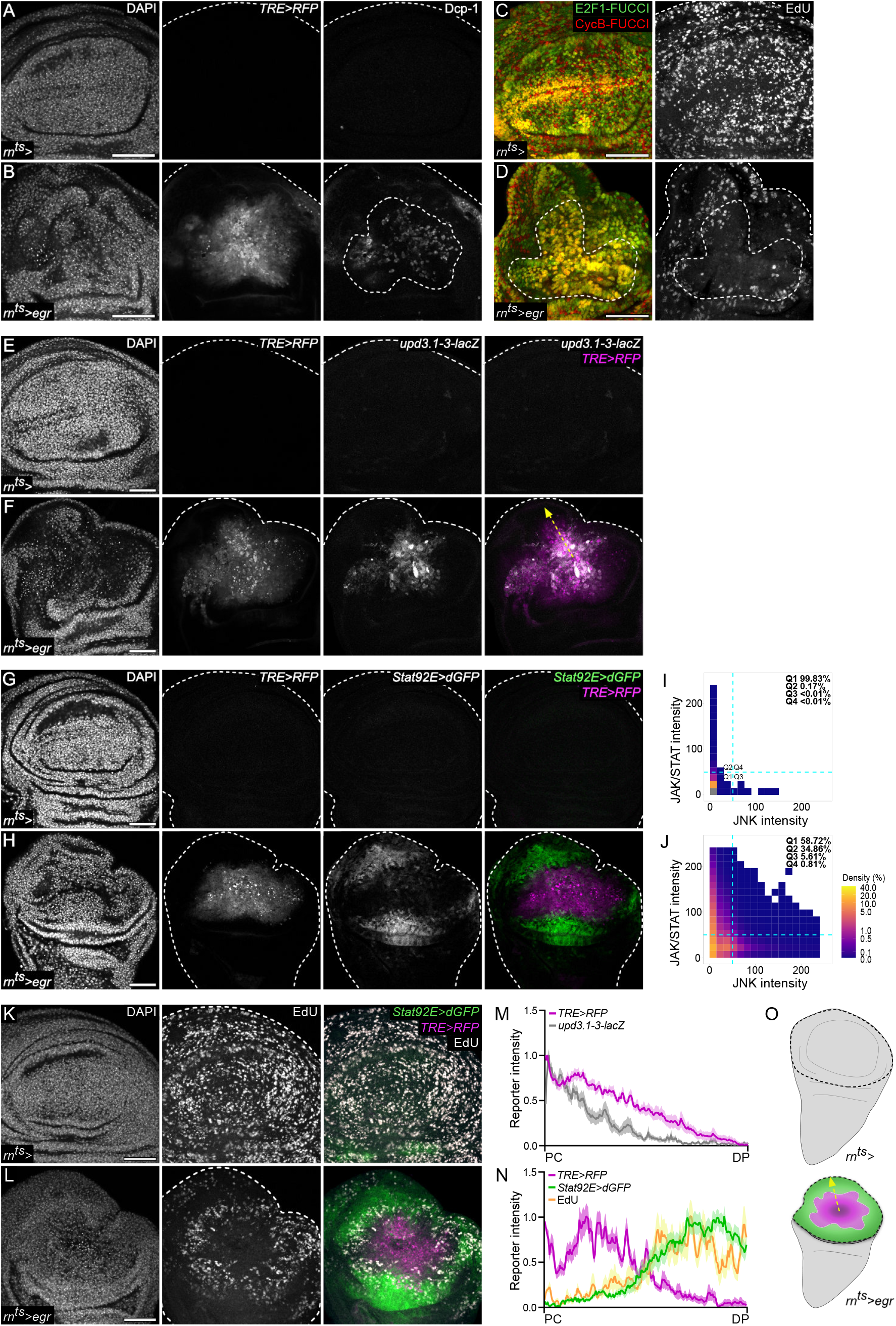
JNK/AP-1 and JAK/STAT are activated in distinct spatial domains upon tissue damage. **(A-B)** A control wing disc **(A)** and wing disc after 24h of *egr*-expression **(B)** in the pouch (R0) (see also **S1.1, B**) under the control of the inducible *rn-GAL4* (*rotund-GAL4*) driver (*rn^ts^>*) in the pouch domain. Discs express the JNK/AP-1 reporter *TRE>RFP* and were stained for cleaved Dcp-1 to visualize apoptosis. **(C-D)** A control wing disc **(C)** and wing disc after 24h of *egr*-expression **(D)** in the pouch (R0), under the control of the inducible *rn-GAL4* (*rotund-GAL4*) driver (*rn^ts^>*). Discs also express the FUCCI cell cycle reporters *mRFP-NLS-CycB^1-266^* (red) and *GFP-E2f1^1-230^* (green) (see also **S1.1A**) and were assessed for DNA replication by EdU incorporation assay. **(E-F)** A control **(E)** and *egr*-expressing disc **(F)** at R0, expressing the *upd3.1-3-LacZ* (gray) and the JNK/AP-1 reporter *TRE>RFP* (magenta). Discs were stained for β-Galactosidase. **(G-H)** A control **(G)** and *egr*-expressing disc **(F)** at R0, expressing the JNK/AP-1 reporter *TRE>RFP* (magenta) and the dynamic JAK/STAT reporter *Stat92E>dGFP* (green). **(I-J)** 2D density plots mapping pixel fluorescence intensity values for the *TRE>RFP* (JNK intensity, X-axis) and *Stat92E>dGFP* (JAK/STAT intensity, Y-axis) reporters, measured in the combined wing pouch and hinge domains (black dashed lines in **O**) in control, *rn-GAL4*-expressing discs **(I)** or *egr*-expressing discs **(J)**. Dashed lines (cyan) represent visually defined thresholds distinguishing low or high reporter activity in the quadrants Q1,Q2,Q3 and Q4. Note that less than 1% of pixels show both high JNK and high JAK/STAT signaling activity in the combined JNK and JAK/STAT signaling domains (i.e. pouch and hinge). Graphs represent pooled values from *n=4*, control and *n=3, egr*-expressing discs. **(K-L)** A control **(K)** and *egr*-expressing disc **(L)** assessed for S-phase activity using EdU incorporation assays. Discs also express the *TRE>RFP* (magenta) and *Stat92E>dGFP* (green) reporters. **(M-N)** Fluorescence intensity profiles for *upd3.1-3LacZ* (gray) and *TRE>RFP* (magenta) reporters **(M)**, or *TRE>RFP* (magenta), *Stat92E>dGFP* (green) reporters and EdU (yellow) reporter intensities **(N)** traced from the pouch center (PC) to the disc periphery (DP) (yellow arrows in **F** and **O**). Graph represents mean ± SEM of reporter fluorescence intensity values from *n=15* tracks across a representative disc, scaled to the maximum measured value. **(O)** Schematic representation of undamaged control discs and spatially segregated patterns of JNK/AP-1 (magenta) and JAK/STAT (green) reporter activity in 24h *egr*-expressing discs (R0). Maximum projections of multiple confocal sections are shown in **K-L**. Discs were stained with DAPI to visualize nuclei. Scale bars: 50 µm

Upon tissue damage, *upd’s* - the pro-mitogenic cytokines for the JAK/STAT pathway - are generally upregulated at wound sites where JNK/AP-1 activity is high (Fig. 1E,F,M, see also S1.1F,G) [20, 28, 49, 51]. Consistent with their paracrine function [67, 68], JAK/STAT signaling was broadly induced in the pouch periphery and hinge of *egr-*expressing discs (Fig. 1G,H), coinciding with regions of proliferation (Fig. 1K,L,N). Yet, high JNK-signaling cells in *egr-*expressing discs that upregulated the transcriptional reporters for *upd1* and *upd3* (Fig. S1.1F,G; Fig. 1E,F,M) [69] did not undergo proliferation and, importantly, did not exhibit JAK/STAT activity, as assessed by the *10xStat92E>dGFP reporter* (Fig. 1I,J) [70]. This was in stark contrast to wing disc development where ligand expression and pathway activation patterns largely coincide [70–73]. Our unexpected finding that JAK/STAT signaling is low in *upd*-producing cells, strongly suggests that a distinct regulatory network controls JAK/STAT activation during JNK-driven responses to tissue damage.

To further support this conclusion, we first wanted to rule out that the observed JAK/STAT activation pattern in the periphery of *egr-*expressing discs perpetuated from early hinge patterns of developing discs due to a Dilp8-induced developmental delay (Fig. S1.2 A) [74]. Hence, we monitored temporal changes in JAK/STAT activity via a time course of 0h, 7h and 14h of *egr-*expression. We consistently observed ectopic induction of the JAK/STAT reporter in the hinge, above developmental levels and with an altered pattern (Fig. S1.2 B-M). This data confirmed that *upd1-3*-driven JAK/STAT activity is induced *de novo* in the hinge and pouch periphery upon tissue damage.

In summary, we find that JNK/AP-1 activation in the *egr-*expressing pouch coincides with the expression of *upd* cytokines. We show that *upd* cytokines activate JAK/STAT only non-autonomously in a cell population spatially segregated from high JNK-signaling and *upd*-producing cells. In addition to the well-described non-autonomous activation of JAK/STAT by JNK/AP-1 activation, this strongly indicates that cell-autonomous repression of JAK/STAT signaling may be an important consequence of JNK/AP-1 activation (Fig. 1O). This is consistent with recent observations of JNK/AP-1 and JAK/STAT activity segregation during tissue stress in other tissue types, such as epidermal wounds [14]. We therefore demonstrate a recurring theme of spatial segregation of JNK/AP-1 and JAK/STAT signaling upon tissue damage. Importantly, we also show that such spatial patterns of segregating JNK/AP-1 and JAK/STAT signals strongly correlate with spatial segregation of regenerative cell behaviors, specifically G2 stalling and proliferation, into spatially distinct signaling domains.

### Figure 2 JNK/AP-1 signaling represses JAK/STAT activity

We wanted to first understand whether JNK/AP-1 directly represses JAK/STAT activity and thereby restricts it to the wound periphery. We therefore tested if clonal expression of JNK/AP-1 directly represses JAK/STAT activity, by co-expressing a constitutively activated form of the JNKK Hep along with p35 to prevent cell death [75, 76]. Indeed, *hep^act^* clones cell-autonomously repressed JAK/STAT activity (Fig. 2A-D, S2A), both when non-autonomously activating it in surrounding cells in the peripheral pouch, hinge or peripodium (Fig. 2A-C, S2A), as well as when within developmentally patterned activity in the hinge (Fig. 2D) [72]. These experiments demonstrate that JNK/AP-1 represses JAK/STAT signaling cell-autonomously, even when JNK/AP-1 has the ability to activate JAK/STAT non-autonomously.

**Figure 2.**
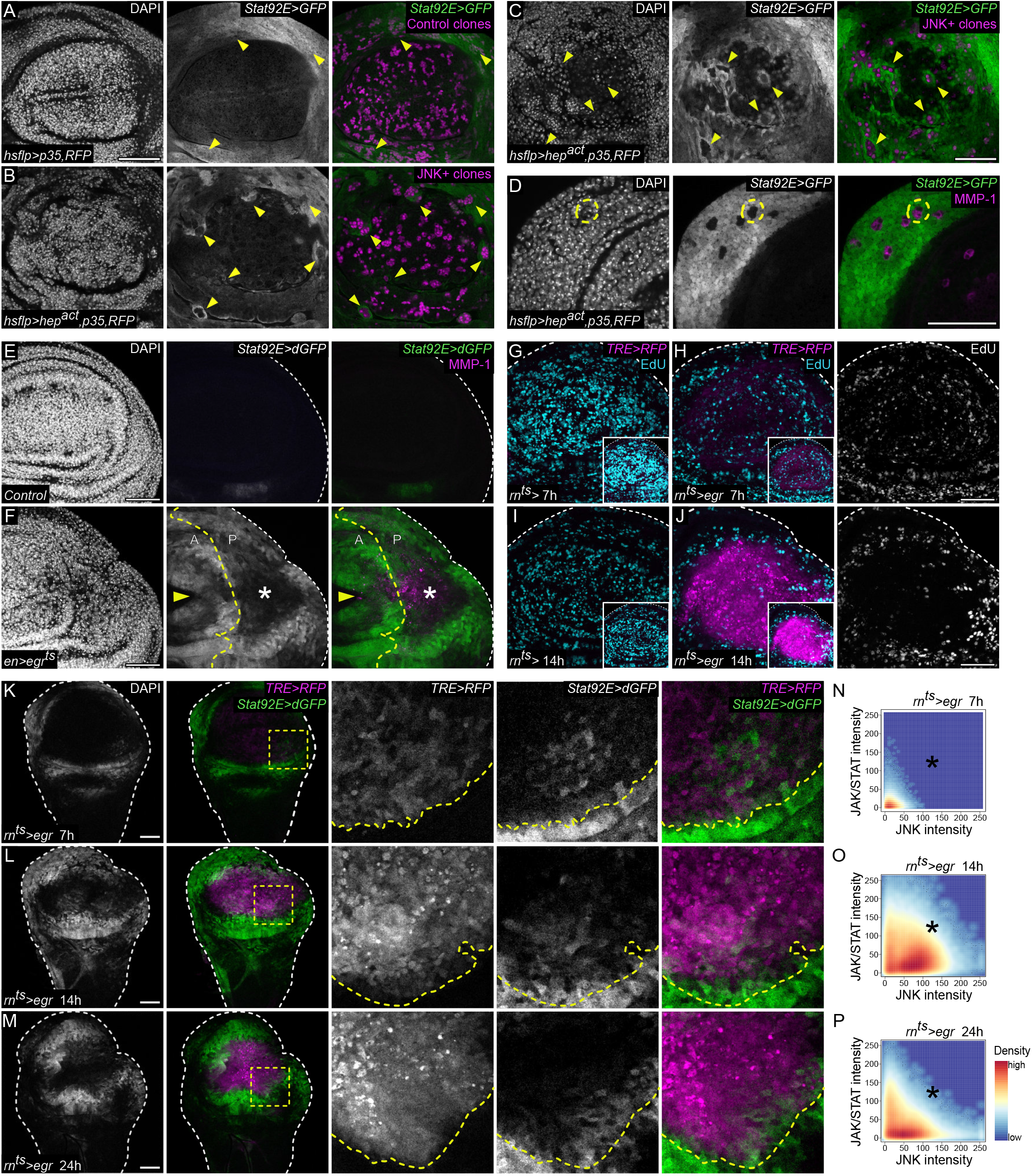
JNK/AP-1 signaling represses JAK/STAT activity cell-autonomously. **(A-C)** RFP expression marks *p35*-expressing control clones in the pouch (magenta, **A**) or *p35,hep^act^*-expressing JNK+ clones in the pouch (magenta, **B**) and peripodium (magenta, **C**) 28h after clone induction in discs expressing the stable *Stat92E>GFP* reporter (green). Yellow arrowheads indicate clones of interest. Note the cell-autonomous repression of *Stat92E>GFP* reporter activity in JNK+ clones in the pouch periphery and hinge **(B)**, as well as in the peripodium **(C)**, when compared to control clones in the pouch **(A)** and peripodium (see **S2A**) **(D)** *p35,hep^act^*-expressing JNK+ clones in a disc expressing the *Stat92E>GFP* reporter (green), stained with anti-MMP-1 (magenta) - a downstream target of the JNK/AP-1 signaling pathway. The yellow circle highlights a JNK+ clone of interest, which shows a strong cell-autonomous repression of the JAK/STAT reporter activity in the JAK/STAT-signaling competent hinge, 48h after clone induction. **(E-F)** Control disc **(E)** and a disc expressing *egr* **(F)** in the posterior compartment using the *enGAL4* driver at R0. Dashed yellow line marks the A/P boundary. Both discs express the *Stat92E>dGFP* reporter (green) and are stained for the JNK/AP-1 target MMP-1 (magenta). White asterisk indicates JNK/AP-1 reporter activity in the posterior pouch region. Note the non-cell autonomous JAK/STAT reporter activity in the anterior pouch, indicated by yellow arrowheads in **F**. **(G-J)** Control **(G,I)** and *egr*-expressing **(H,J)** discs at 7h or 14h after ablation expressing the *TRE>RFP* reporter (magenta) and assessed for S-phase activity using EdU incorporation assay (cyan). While the number of S-phase cells are uniformly distributed in the pouch domain of undamaged control discs **(G,I)**, note the successive decrease in the S-phase cells (cyan) from 7h **(H)** to 14h **(J)** of *egr*-expression, coinciding with increasing *TRE>RFP* reporter activity in the pouch. Insets show the same images, with settings adjusted to visualize the JNK/AP-1 activity in the pouch of 7h *egr*-expressing discs. **(K-M)** A time-course analysis of *TRE>RFP* (magenta) and *Stat92E>dGFP* (green) reporter activity in *egr*-expressing discs after 7h **(K)**, 14h **(L)** and 24h **(M)** of ablation. Yellow squares show magnified regions. Yellow lines mark the interface between JNK/AP-1 and JAK/STAT signaling cells. Note the increasing dominance of JNK/AP-1 signaling and decreasing JAK/STAT activation inside the boundary of JNK/AP-1 signaling cells at the JNK/AP-1-JAK/STAT interface. **(N-P)** Representative 2D density plots of pixel fluorescence intensity for the *TRE>RFP* (JNK intensity, X-axis) and *Stat92E>dGFP* (JAK/STAT intensity, Y-axis) reporters, measured exclusively within the JNK/AP-1 signaling pouch domain (magenta in Fig. 1O) from 7h **(L)**, 14h **(M)** and 24h **(N)** *egr*-expressing discs. Black asterisks highlight the region of increase and subsequent decline in the density of co-localizing pixels with medium and high reporter activities from 7h to 14h **(N,O)**, and 14h to 24h **(O,P)** respectively. Maximum projections of multiple confocal sections are shown in **E-J**. Discs were stained with DAPI to visualize nuclei. Scale bars: 50 µm

In these mosaic experiments, hinge cells activated JAK/STAT signaling more robustly than cells of the central pouch (Fig. 2A-B), likely as a result of developmental pre-patterns conferring distinct competence [72]. Namely, the transcription factors Nubbin (Nub) and Rotund (Rn) reduce developmental Stat92E activity in the pouch [77]. To first exclude the possibility that JNK/AP-1 represses JAK/STAT by inducing pouch-specific transcription factors [77], we assessed Nub and Rn levels within *hep^act^* clones. *hep^act^* clones did not ectopically induce Nub or Rn, in fact, Nub was distinctly downregulated (Fig. S2B-D). Furthermore, we expressed *egr* in the posterior compartment of the wing disc and examined whether JAK/STAT could be activated in the adjacent anterior pouch (Fig. S1.1B). The cells highly positive for the JNK/AP-1 target gene - MMP-1 [78] did not activate JAK/STAT signaling. However, JAK/STAT signaling could be distinctly activated in the surrounding tissue, including the anterior pouch (Fig. 2E,F). These experiments confirm that JAK/STAT signaling can *in principle* be activated *de novo* in the pouch by cell non-autonomous activity of JNK/AP-1 signaling cells. Altogether, these experiments demonstrate that tissue damage associated with high levels of JNK/AP-1 signaling repress JAK/STAT activity in a cell autonomous manner, independent of developmental competence.

Accordingly, we wanted to better understand if JAK/STAT repression by JNK/AP-1 may depend on JNK/AP-1 signaling strength or time. We therefore analyzed the temporal evolution of JNK/AP-1 and JAK/STAT activity in the JNK-signaling domain of *egr-*expressing discs. A time course analysis revealed that JAK/STAT was mildly elevated during short-term JNK-signaling in the pouch by 7 h of *egr-*expression, and increased even further with 14 h of *egr-*expression (Fig. 2K,L,N,O, Fig. S2E,F,H,I,K,). Interestingly, cells displayed co-existence of both pathways at moderate levels, but also often displayed exclusive activation of either reporter (Fig. S2L,M). After 24 h of sustained *egr-*expression, high JNK/AP-1 signaling was observed in the pouch, but notably, JAK/STAT activity had now largely disappeared from this domain (Fig. 2M,P, Fig. S2G,J,K). These observations reveal that short-term or moderate activity of JNK/AP-1 facilitates local, intermixed activation of JAK/STAT. However, sustained and high activation of JNK/AP-1 efficiently excludes JAK/STAT activity cell autonomously. As a consequence, JNK/AP-1 and JAK/STAT signaling are increasingly segregated into distinct cell populations within the tissue.

Notably, these signaling dynamics are synchronously followed by cell behavior dynamics. We observed that prolonged JNK/AP-1 signaling correlated with an increased exclusion of proliferation from the pouch (Fig. 2G-J; Fig. 1K,L,N). Altogether, these data suggest that chronic and high levels of JNK/AP-1 activation drive spatial wound site signaling and cell behavior patterns from a local intermixed state to a larger, segregated and bistable field.

### Figure 3 JNK/AP-1 acts as a wound organizer by employing a mutual-repression network

We wanted to understand the type of regulatory interactions between JNK/AP-1 and JAK/STAT pathways that establish spatially segregated signaling domains on a tissue scale [79–82]. Regulatory networks describe how components of different signaling pathways interact [83–85] or are integrated to define a cell’s signaling state and establish distinct cellular outcomes [86, 87]. How regulatory networks can also organize cellular behaviors to influence tissue-level outcomes has been extensively described using mathematical modeling [88–90].

To characterize the rules of the regulatory network between JNK/AP-1 and JAK/STAT that ultimately drives emergence of spatial segregation to organize tissue stress responses, we developed a mathematical model using partial differential equations. These described the spatio-temporal evolution of JNK/AP-1 and JAK/STAT activation depending on relative concentrations of components in the system (Fig. 3A-C) [86, 91]. We defined a set of experimentally determined rules, namely: (1) JNK/AP-1 effectors propagate JNK/AP-1 via production of paracrine factors, such as *egr* or diffusible ROS [28, 54, 92]; (2) JNK/AP-1 effectors activate JAK/STAT via production of paracrine factors, such as *upd*’s [20, 28, 49, 51]; (3) Both pathways use positive feedback loops to enhance and stabilize their own activation [42, 55, 56, 93] and (4) JAK/STAT represses JNK/AP-1 cell-autonomously, via the ZEB-1 homologue Zfh2 (Fig. S3A-C) [51]. To simplify the model (Fig. 3B), we excluded negative feedback loops on JNK/AP-1 and JAK/STAT [94–96], as they often have a more modulatory function in noise buffering, temporal processing or saturation dynamics [97].

**Figure 3.**
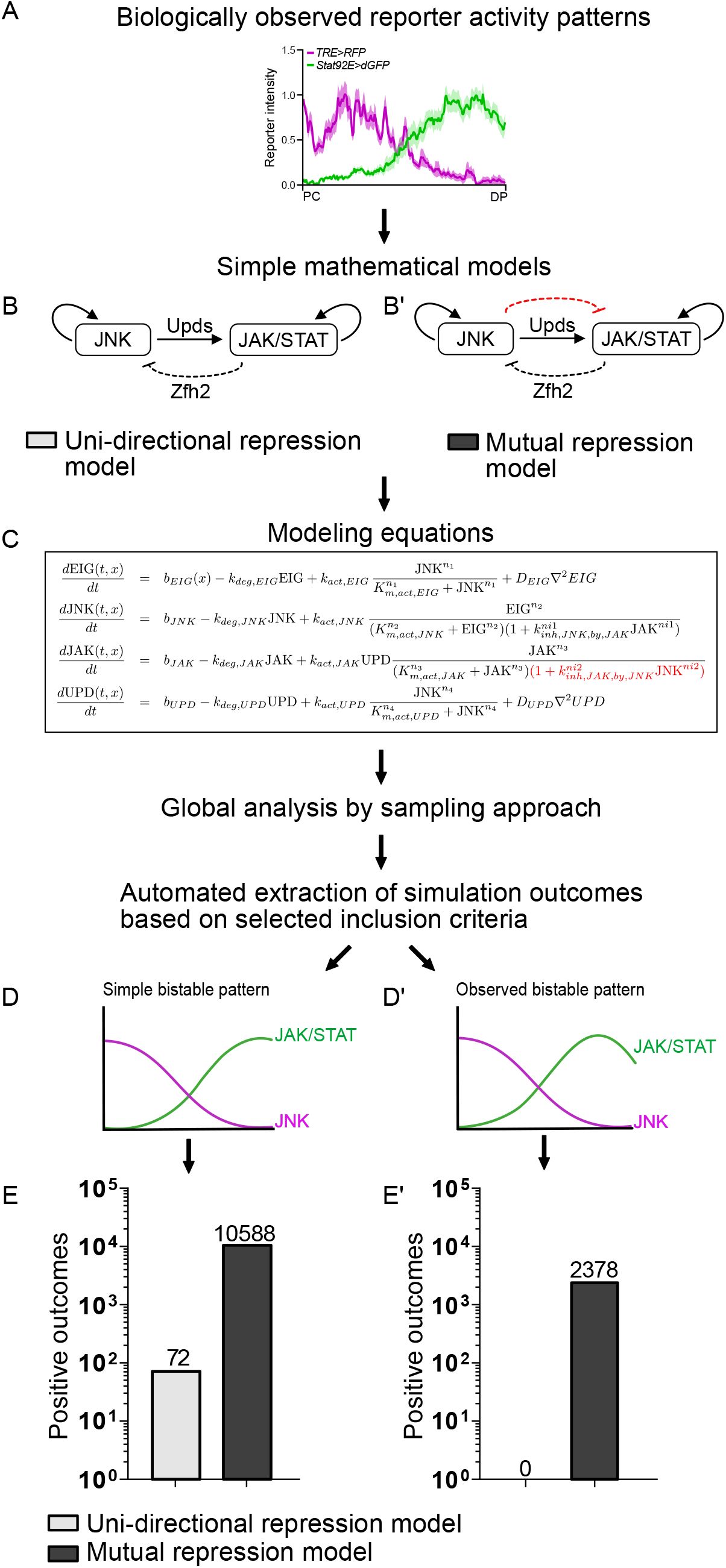
A mutual-repression loop promotes bistable segregation of JNK/AP-1 and JAK/STAT activation patterns. **(A-E’)** Workflow to investigate the presence of a regulatory motif between the JNK/AP-1 and JAK/STAT signaling pathways driving pattern formation in *egr*-expressing discs. Fluorescence intensity profiles for the *TRE>RFP* (magenta) and *Stat92E>dGFP* (green) reporter patterns show bistable segregation in 24h egr-expressing discs. Intensity profiles are traced from the pouch center (PC) to the disc periphery (DP) **(A)**. Schematic showing the models tested – a uni-directional repression model acting only on JNK/AP-1 signaling **(B)** and a mutual repression model acting on both JNK/AP-1 and JAK/STAT signaling **(B’)**. Modeling equations were generated, and a global analysis by sampling approach was used to analyze 21 selected parameters **(C)** (see also Fig.**S3**) and test for outcomes that produce bistable, switch-like patterns **(D-D’)**. Number of simple bistable outcomes **(D)** from the mutual repression model (dark grey) are substantially greater than those from the uni-directional repression model (light grey) **(E)**. Applying stringent selection criteria based on the biologically observed reporter activity pattern **(D’)**, positive outcomes reduce to 0 in the model without the mutual repression **(E’)**.

As we found experimentally that JNK/AP-1 repressed JAK/STAT cell-autonomously, we also designed a mutual repression model that, together with the aforementioned rules, included a rule that JNK/AP-1 represses JAK/STAT signaling (Fig. 3A,B’) [98]. Using an unbiased sampling approach of a large parameter space, we compared the ability of the two models to produce simple bistable patterns (Fig. 3D). We found that the mutual repression model produced 140-fold more positive outcomes than a uni-directional model (10588 vs. 72 solutions) (Fig. 3E). When we switched from scoring simple bistable patterns (Fig. 3D) to observed bistable patterns (Fig. 3D’) of JNK/AP-1 and JAK/STAT signaling in *egr*-expressing wing discs, the difference in competencies between the two models became even more prominent (Fig. 3E’). The uni-directional model was not sufficient to recapitulate observed bistable patterns, which suggests that the rule of JNK/AP-1 cell-autonomously repressing JAK/STAT is necessary for generating the segregated JNK/AP-1 and JAK/STAT signaling patterns seen during tissue stress. Accordingly, we show that a mutual-repression network lies at the core of JNK/AP-1 and JAK/STAT signaling interactions in response to tissue damage. Supporting this result, mutual repression networks have been extensively described to guide the formation of bistable states [99–105]. Furthermore, these networks can be further enhanced by self-activation loops, such as those integrated in our model [98, 102, 106]. Indeed, if JNK/AP-1 and JAK/STAT cross-talk were to completely lack mutual repression, JNK/AP-1 and JAK/STAT signaling would be expected to expand from the wound site in an unrestrained manner. Thus, the mutual repression regulatory network also ensures that these signaling pathways remain restricted during regeneration.

The length scale-independent space utilized by the model raises the possibility that this mutual repression network could act at length scales ranging from neighboring cells to multicellular tissues. *In vivo*, length scales may depend on the level of network activation and thus the concentration of non-autonomously acting cytokines (Egr, Upd’s), as suggested by experiments shown in (Fig. 2K-P and Fig. S1.2). Indeed, we observed that the number of segregating solutions produced by the model increased with increasing rates of JNK/AP-1 activity (Fig. S3D). This observation parallels the progressive segregation of signaling domains found experimentally upon chronic activation of JNK/AP-1 signaling (Fig. 2K-P).

Taken together we propose that JNK/AP-1 initiates a mutual repression network between JNK/AP-1 and JAK/STAT signaling that, via the production of Upd’s, facilitates the spatial segregation of these two pathways. The spatial segregation of JNK/AP-1 and JAK/STAT via mutual repression allows the tissue to maintain a JNK-dependent G2 stall in the presence of pro-mitogenic signals, and restricts proliferation to a separate cell population. Thus, induction of JNK/AP-1 signaling upon tissue damage and mutual repression with JAK/STAT is sufficient to set up temporal and spatial signaling patterns, and functionally distinct cell populations.

### Figure 4 Ptp61F represses JAK/STAT activity in JNK/AP-1 signaling cells

To understand how JNK/AP-1 may cell-autonomously repress JAK/STAT activation, we first asked if JNK/AP-1 downregulates core components of the JAK/STAT pathway. Our previous study demonstrated that the core components *dome*, *hop* and *stat92E* [107] were not transcriptionally altered in *egr-*expressing discs [51]. Supporting this observation, we found that the expression of GFP-tagged Hop [108] and Stat92E proteins from functionally validated lines were unaltered in *egr-*expressing cells (Fig. S4.1A-I). This suggested that repression of JAK/STAT signaling was likely due to altered function of regulators of JAK/STAT signaling. We thus analyzed known negative regulators in *egr-*expressing discs. Levels of dPIAS/Su(var)2-10 [109] and Apontic [110] were unchanged, but *ken* expression [111] was elevated in *egr-*expressing discs (Fig. S4.2A-F). However, neither knock-down of *ken* nor *dPIAS/Su(var)2-10* increased JAK/STAT activity in *egr-*expressing cells (Fig. S4.2G-L). Similarly, we could not reproduce findings that knockdown of *apontic* reinstated JAK/STAT signaling in *egr-*expressing discs (Fig. S4.2M-O) [112].

We then turned to the tyrosine phosphatase *Ptp61F,* a known repressor of Stat92E activation [113, 114] which is transcriptionally induced in JNK-dependent tumors and other stress conditions [23, 115]. Repression of JAK/STAT signaling by Ptp61F may occur at the level of receptor complex signaling or Stat92E activity [113, 114, 116]. We found that knock-down of *ptp61F* caused upregulation of the Stat92E reporter activity in high JNK-signaling cells of *egr-*expressing discs (Fig. 4A-E, Fig. S4.3A-C). This suggests that Ptp61F is specifically required in JNK-signaling cells to suppress Stat92E activity. Hence, to test if dephosphorylation of Stat92E in JNK-signaling cells is rate-limiting for JAK/STAT signaling, we overexpressed an HA-tagged Stat92E [117] in *egr-*expressing cells to saturate the dephosphorylation capacity of Ptp61F. Indeed, *egr,stat92E-HA*-coexpressing cells in the central pouch domain activated the JAK/STAT reporter at higher levels when compared to *egr-*expressing discs (Fig. 4F-J, Fig. S4.3D,E). In contrast, co-expression of the JAK *hop* failed to induce JAK/STAT reporter activity in *egr-*expressing discs (Fig. S4.3F-I). Importantly, this demonstrates that while signal transduction from Upd ligands down to the effector Stat92E is in principle active in JNK-signaling cells, it is the Ptp61F-mediated dephosphorylation of Stat92E that becomes rate-limiting for JAK/STAT signaling.

**Figure 4.**
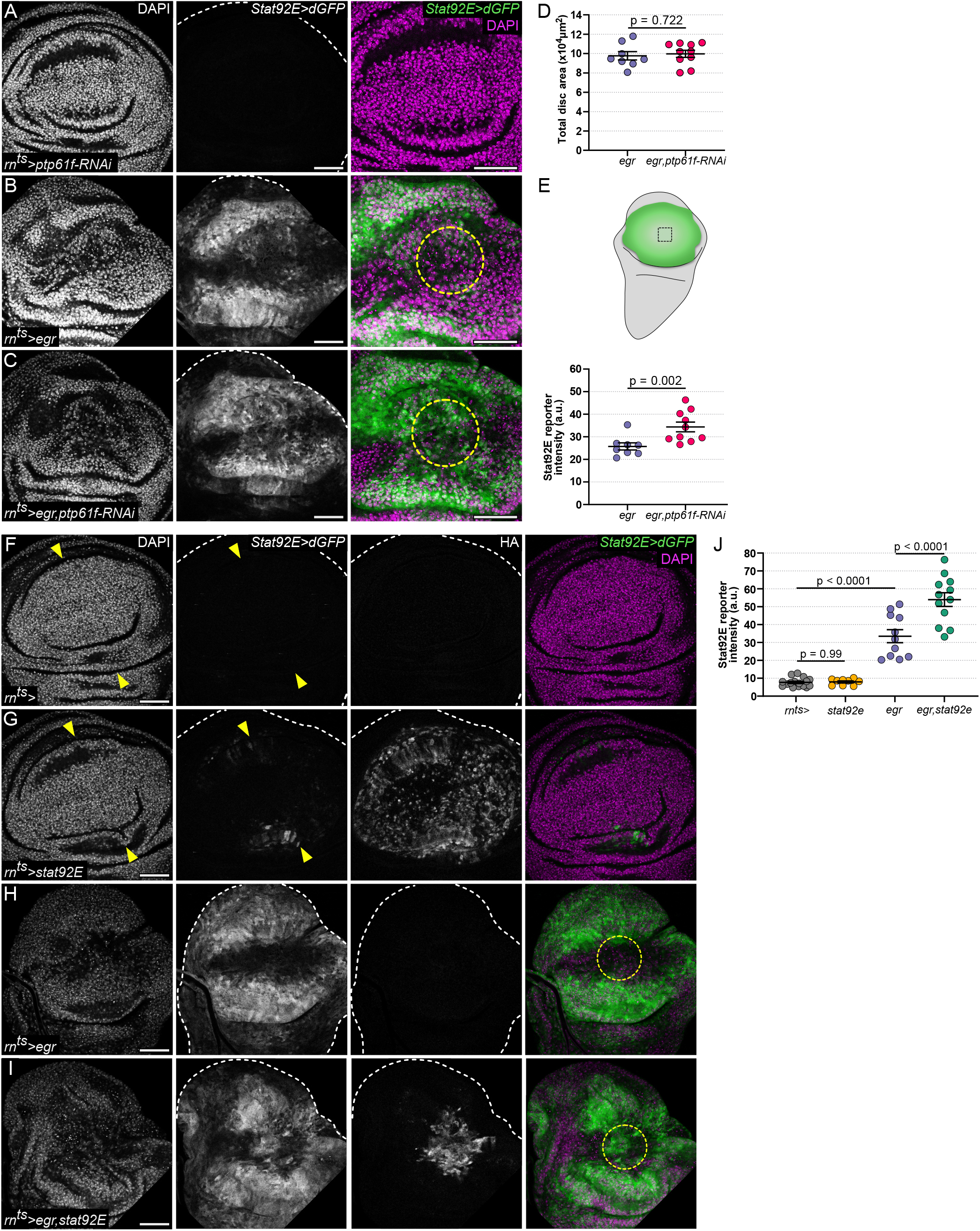
Ptp61F represses JAK/STAT activity in JNK/AP-1 signaling cells. **(A-C)** A *ptp61F-RNAi*-expressing **(A)**, *egr*-expressing **(B)** and *egr,ptp61F-RNAi*-co-expressing disc **(C)** at R0. All discs also express the *Stat92E>dGFP* reporter (green). Yellow circles highlight the central domain of remaining *rn-GAL4* expressing cells. Note the derepression and ectopic JAK/STAT activity in the *egr,ptp61F-RNAi*-co-expressing disc. **(D)** Quantification of total area of wing discs. Graphs display mean ± SEM for *n=8*, *egr*-expressing and *n=10*, *egr,ptp61F-RNAi*-co-expressing discs. T-tests were performed to test for statistical significance. **(E)** Quantification of the *Stat92E>dGFP* reporter fluorescence intensity measured within the JNK/AP-1 signaling pouch domain. Black square in schematic shows measured region. Graphs display mean ± SEM for *n=8*, *egr*-expressing and *n=10*, *egr,ptp61F-RNAi*-co-expressing discs. U-tests were performed to test for statistical significance. **(F-I)** A control **(F)**, *stat92E*-expressing **(G)**, *egr*-expressing **(H)** and *egr,stat92E*-co-expressing disc **(I)**. All discs also express the dynamic *Stat92E>dGFP* reporter (green). Staining for HA confirms expression and nuclear/cytoplasmic localization of the *UAS-stat92E-3xHA* construct. Yellow arrowheads point to the restricted, non-autonomously induced JAK/STAT activity in the peripheral pouch in control discs **(F,G)**. Yellow circles highlight the central domain of remaining *rn-GAL4* expressing cells **(H,I)**. **(J)** Quantification of the *Stat92E>dGFP* reporter fluorescence intensity measured within the JNK/AP-1 signaling pouch domain. Black square in schematic **(E)** shows measured region. Graphs display mean ± SEM for *n=16*, control discs (*rn^ts^>*), *n=9*, *stat92E*-expressing discs, *n=11*, *egr*-expressing and *n=12*, *egr,stat92E*-co-expressing discs. One-way ANOVA with multiple comparisons was performed to test for statistical significance. Discs were stained with DAPI (magenta) to visualize nuclei. Scale bars: 50 µm

### Figure 5 JAK/STAT repression is required for the G2 stall and protects central wound cells from apoptosis

As spatial segregation of JNK/AP-1 and JAK/STAT signaling domains was a robust feature of *egr-*expressing discs, we wondered if the mutual repression network was necessary for regeneration. We therefore forced expression of Stat92E in high JNK/AP-1 signaling cells, and closely monitored cell and tissue level responses, such as cell survival and proliferation. The loss of JNK/AP-1 and JAK/STAT segregation led to a pronounced increase in apoptosis (Fig. 5A-D). Co-expression of UAS-GFP confirmed that these apoptotic cells originated from cells where JNK/AP-1 and JAK/STAT were forcibly co-activated (Fig. S5A-D).

**Figure 5.**
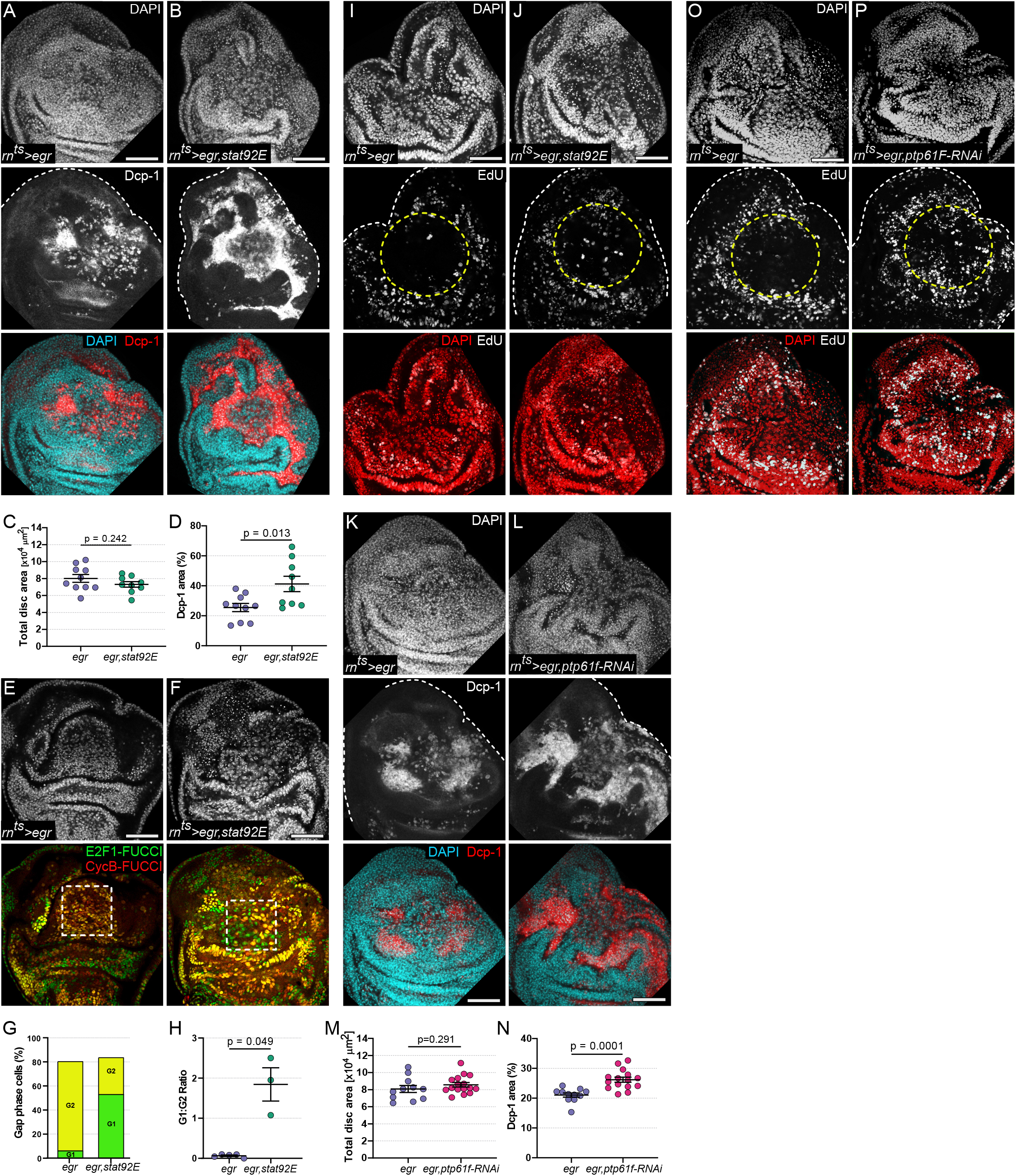
JAK/STAT repression by JNK/AP-1 is required for the G2 arrest and protects central wound cells from apoptosis. **(A-B)** An *egr*-expressing **(A)** and *egr,stat92E*-coexpressing disc **(B)** at R0, stained for DAPI (cyan), and cleaved Dcp-1 (red) to visualize apoptosis. **(C-D)** Quantification of total wing disc area **(C)** and normalized area occupied by cleaved Dcp-1 **(D)**. Graphs display mean ± SEM for *n=10, egr*-expressing and *n=9, egr,stat92E-*co-expressing discs. T-tests were performed to test for statistical significance. **(E-F)** An *egr*-expressing **(E)** and *egr,stat92E*-coexpressing disc **(F)** at R0, also expressing the FUCCI reporter. Dashed white squares highlight the distinct shift from G2 (yellow cells) to G1 (green cells) in the pouch domain. **(G-H)** Bar graph **(G)** representing the cumulative proportion of gap phase cells in G1 (green) and G2 (yellow) for each genotype. Graph **(H)** displays G1-phase:G2-phase ratios obtained from *n=5, egr*-expressing and *n=3, egr,stat92E-*co-expressing discs. Dashed white squares in **E-F** show measured regions. T-test with Welch’s correction was performed to test for statistical significance. **(I-J)** EdU incorporation assay to detect S-phase cells (gray) in *egr*-expressing **(I)**, or *egr,stat92E*-co-expressing **(J)** disc. Discs were stained with DAPI (red) to visualize nuclei. **(K-L)** An *egr*-expressing **(K)** and *egr,ptp61F-RNAi*-coexpressing **(L)** disc at R0, stained for DAPI (cyan), and cleaved Dcp-1 (red) to visualize apoptosis. **(M-N)** Quantification of total wing disc area **(M)** and normalized area occupied by cleaved Dcp-1 **(N)**. Graphs display mean ± SEM for *n=11, egr*-expressing and *n=15, egr,ptp61F-RNAi-*coexpressing discs. T-tests were performed to test for statistical significance. **(O-P)** EdU incorporation assay to detect S-phase cells (gray) in an *egr*-expressing **(O)**, and *egr,ptp61F-RNAi*-coexpressing **(P)** disc. Discs were also stained with DAPI (red) to visualize nuclei. Maximum projections of multiple confocal sections are shown in **A-B, I-L, O-P**. Scale bars: 50 µm

We thus hypothesized that JAK/STAT interfered with the protective G2-stall of high JNK-signaling cells, and thereby increased apoptosis [31, 37], While Stat92E expression altered the proportion of S-phase cells in undamaged control discs, we observed no changes in the proportion of cells in G1 or G2 (Fig. S5E-H). In contrast, a cell cycle analysis of *egr,Stat92E-HA* co-expressing domains revealed a substantial increase in G1 and S-phase cells, indicating that JNK-signaling cells failed to stall in G2 (Fig. 5E-J). Thus, forced Stat92E activation in high JNK-signaling cells overrides the protective JNK-induced G2 stall and consequently increases apoptosis. Supporting these results, we found that knock-down of Ptp61F in *egr-*expressing discs also led to an increase in cycling cells in the tissue (Fig. 5O,P) correlating with increased levels of apoptosis (Fig. 5K-N). This highlights the necessity of JNK/AP-1 and JAK/STAT signaling segregation within the tissue, as it maintains the G2 stalled cell population. Thus, the JNK-induced mutual repression network ensures the establishment of two distinct and indispensable cell populations - one that stalls in G2 to prevent apoptosis and secrete necessary pro-mitogenic factors like Upd and other paracrine effectors like Dilp8 and ImpL2 [20, 74], and a second that is able to respond to mitogenic signals and undergoes proliferation necessary for regeneration. The strong similarities with paracrine signaling centers regulating spatial patterning networks linked to specific cell behaviors and fate during development encourages us to propose that the JNK-induced mutual repression network with JAK/STAT functions as a central wound organizer network [118, 119].

### Figure 6 The wound organizer network segregates JNK/AP-1 and JAK/STAT signaling to drive oncogenic growth

Our observation that tissue repair processes after chronic damage support the establishment of a signaling-sending and a responding cell population, which are respectively G2-stalled or proliferative, led us to speculate that such a wound organizer network could well-support a tumor microenvironment. Interestingly, the coexistence of JNK/AP-1 and JAK/STAT pathways in tumors has been extensively described [18–23] and the existence of signal-sending G2-stalled cells controlling non-autonomous proliferation has been demonstrated [31]. However, their spatial patterns have not been well defined.

To understand if JNK/AP-1 and JAK/STAT also engage in a mutual repression network under tumor conditions, we analyzed discs with a reduction in the well-characterized *scrib* tumor-suppressor gene [120] or ectopic expression of oncogenic *Ras^V12^* [121, 122]. To generate a large genetically homogenous tissue, we used the pouch specific *rn-GAL4* driver to express *Ras^V12^* or *scrib-RNAi* in the entire wing pouch. A wing pouch expressing oncogenic *Ras^V12^* for 44 h did not activate JNK/AP-1 or JAK/STAT reporters, demonstrating that Ras^V12^ alone does not co-opt either pathway (Fig. 6A,B, Fig. S6A,B). RNAi-driven knock-down of *scrib* induced barrier dysfunction (Fig. S6C,D) and, consistent with previous reports [123, 124], moderately activated both JNK/AP-1 and JAK/STAT in the pouch (Fig. S6E-H). A closer examination, however, revealed that the *scrib-RNAi* expressing pouch predominantly segregated JNK/AP-1 and JAK/STAT signaling at the level of small cell clusters, while only few cell clusters displayed co-activation of both pathways (Fig. 6C). This pattern mirrored our observations of short-term *egr-*expression (Fig. S2L,M), thus suggesting that different contexts of mild JNK-activation cause segregation at short distances. These observations indicate that JNK/AP-1 activation upon barrier dysfunction via disruption of cell polarity also induces the wound organizer network, which subsequently sets up a bistable segregation of both pathways even at a short range between neighboring cells.

**Figure 6.**
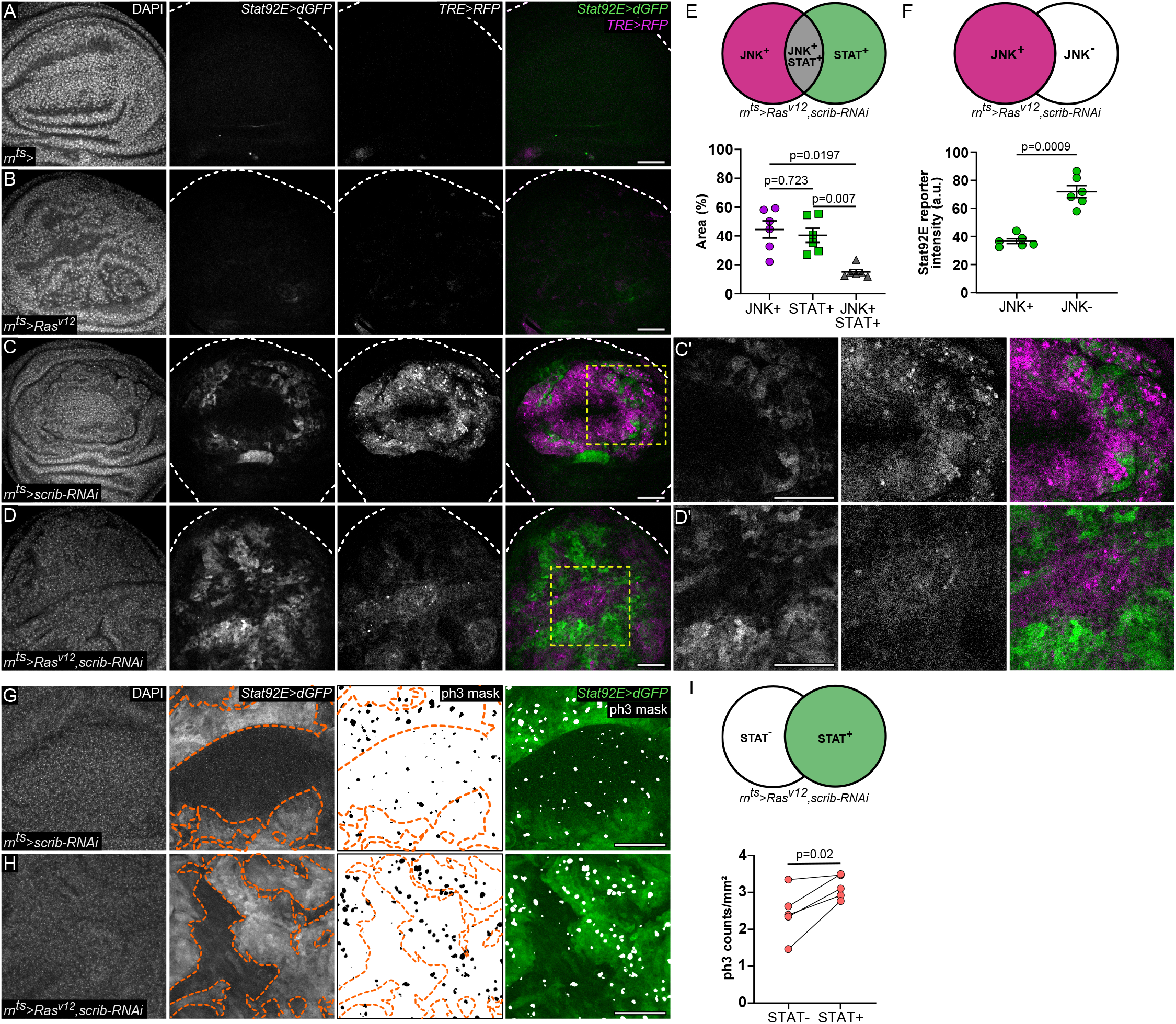
The wound organizer network segregates JNK/AP-1 and JAK/STAT signaling to drive oncogenic growth. **(A-D)** A control wing disc **(A)** and wing discs expressing *Ras^v12^* **(B)**, *scrib-RNAi* **(C, C’)**, or *Ras^v12^*,*scrib-RNAi* **(D,D’)** for 44 h D6 AED. Discs also express the JNK/AP-1 reporter *TRE>RFP* (magenta) and the JAK/STAT reporter *Stat92E>dGFP* (green). Yellow squares in mark area of magnification shown in **(C’,D’)**. **(E)** Schematic shows the *TRE>RFP*-positive (JNK^+^, magenta), *Stat92E>dGFP*-positive (STAT^+^, green) and JNK/AP-1 and JAK/STAT double positive regions (JNK^+^STAT^+^, gray) selected for measurements. Graph represents normalized area of measured regions based on the total signaling area. Note that only 15% of regions within the tumor express both the JNK/AP-1 and JAK/STAT signaling reporters. Graph represents mean ± SEM for *n=6, Ras^v12^,scrib-RNAi*-expressing discs. One-way paired ANOVA with Holm-Šídák’s multiple comparison test was performed to test for statistical significance. **(F)** Schematic shows *TRE>RFP*-positive (JNK^+^, magenta) and *TRE>RFP*-negative (JNK^-^, white) regions selected for measurements. Graph represents *Stat92E>dGFP* fluorescence intensity measured within selected JNK^+^ and JNK^-^ regions. Note the reduction in the reporter intensity from 71.91±4.32 to 36.66±1.74 within the JNK^-^ vs. JNK^+^ region. Graph represents mean ± SEM for *n=6,* ras^v12^,scrib-RNAi-expressing discs. Paired t-tests was performed to test for statistical significance. **(G-H)** A wing disc expressing *scrib-RNAi* **(G)** and *Ras^v12^,scrib-RNAi* **(H)** at 44 h. Discs also express the *Stat92E>dGFP* reporter and were stained for phospho-Histone3 (ph3) to visualize cells in mitosis. Segmented masks of ph3 were generated for quantifications and an overlay of the *Stat92E>dGFP* reporter (green) and ph3 (white) is represented. Orange dashed lines mark the STAT^+^ regions within the discs. **(I)** Schematic shows *Stat92E>dGFP*-positive (STAT^+^, green) and *Stat92E>dGFP*-negative (STAT^-^, white) regions selected for measurements. Graph represents ph3 counts normalized to area in the STAT^+^ and STAT^-^ regions for *n=5, Ras^v12^,scrib-RNAi*-expressing discs. Maximum projections of multiple confocal sections are shown in **G-H** Discs were stained with DAPI to visualize nuclei. Scale bars: 50 µm

Curiously, if Ras^V12^ and *scrib* mutations occur in the same cell, they cooperate to cause dramatic overproliferation of imaginal discs, which is not observed in single *Ras^V12^* or *scrib* mutant tissues. This cooperativity is thought to be driven by activation of JNK/AP-1, as a consequence of *scrib* disrupting cell polarity. JNK/AP-1 signaling then induces *upd’s* and JAK/STAT activation, which can be utilized by a *Ras^V12^* controlled network to drive overgrowth [18, 122]. We thus wondered if Ras^V12^ may disable the mutual repression motif between JNK/AP-1 and JAK/STAT. This would allow autocrine activation of JAK/STAT by JNK signaling in the same cell, as generally described in the literature [18, 122]. Cells driven to cycle despite activation of JNK/AP-1 may then be protected by anti-apoptotic functions of Ras^V12^. We wanted to test this hypothesis and closely monitored JNK/AP-1 and JAK/STAT signaling as well as cell proliferation in Ras^V12^ and *scrib RNAi-*coexpressing discs. Surprisingly, we found that JNK/AP-1 and JAK/STAT activation were still clearly segregated into exclusive signaling domains (Fig. 6D-F, Fig. S6I) with proliferation still predominantly associated with JAK/STAT (Fig. 6G-I). This suggests that even in a Ras^V12^ and *scrib* cooperativity model, where all cells are genetically identical, bistability driven by the mutual repression network organizes the pouch into distinct JNK/AP-1 or JAK/STAT-signaling domains. This observation aligns with previous observations that cooperativity between Ras and *scrib* can rely, in principle, on non-autonomous interactions [18, 125]. Crucially however, our observations demonstrate that non-autonomous cooperation between JNK/AP-1-activating and *Ras^V12^* mutation arises as a self-organizing principle from a wound organizer network creating segregating signaling domains in a genetically homogenous tumorigenic tissue. Moreover, it suggests that activation of this wound organizer network and segregation of signaling and proliferative tasks may provide an advantage to tumors over autocrine integration of JNK/AP-1 and JAK/STAT signaling in the same cell.

## Discussion

Previous studies established a role for JNK/AP-1 in promoting regeneration upon injury [35, 59]. This role is, in part, mediated by the expression of *upd’s* which are essential to promote proliferation and survival via JAK/STAT signaling in a non-autonomous manner [28, 35, 48, 51, 126–129]. By identifying a mutual repression network between JNK/AP-1 and JAK/STAT, we conceptually establish JNK as a core organizer of tissue repair with parallels to organizers of cell fate patterning in developing tissues. The rules of the wound regulatory network we describe have important implications. Previous reports suggest that JNK/AP-1 activates itself non-autonomously, for example via activation of paracrine *egr* or ROS [28, 54, 92]. This implies that JNK/AP-1 activation could result in unchecked spatial expansion of JNK/AP-1-signaling, and accordingly of JAK/STAT signaling. Their unrestrained co-expansion would likely result in chronic inflammation. However, the mutual repression motif restrains the expansion of both JNK/AP-1 and JAK/STAT signaling domains, thus defining stable regions of inflammatory (mitogenic signals) and regenerative (pro-proliferative) responses at the start of the tissue repair process. We demonstrate that molecular mediators of mutual repression are Zfh2 downstream of JAK/STAT [51], and Ptp61F downstream of JNK.

At the level of cell behaviors, JNK/AP-1 and JAK/STAT generate mutually exclusive responses (Fig. 7B). JNK/AP-1 signaling supports wound center behaviors like tissue sealing [24, 25, 78], cell fusion [14], establishment of paracrine signaling and importantly, an apoptosis-resistant state that depends on a G2 cell cycle stall [31]. In contrast, JAK/STAT signaling supports wound-distal behaviors, such as compensatory proliferation, potentially via upregulation of G1-S cyclins such as CycD or CycE [130, 131].

**Figure 7.**
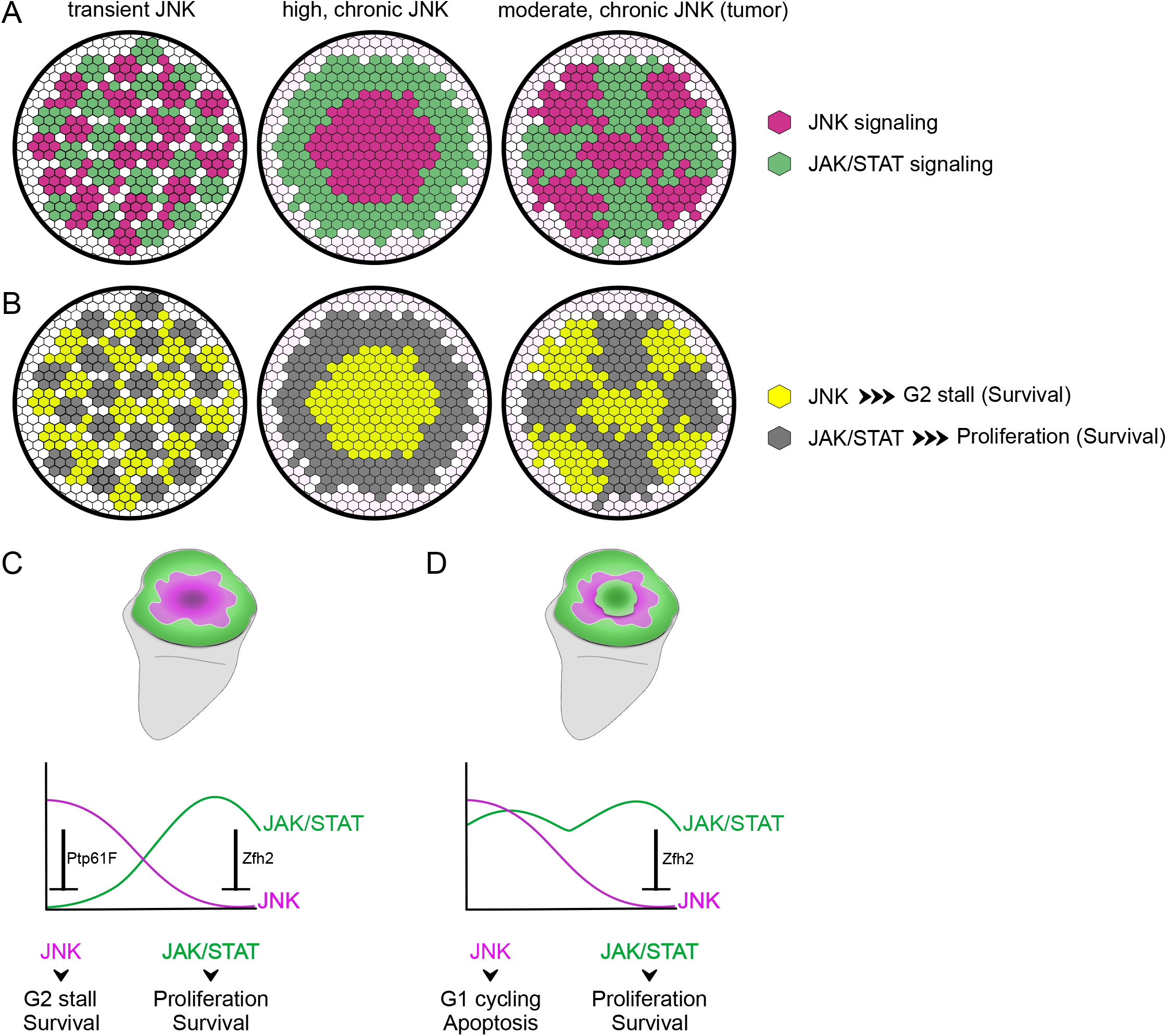
Mutual repression between JNK/AP-1 and JAK/STAT sets up wound organizer activity. **(A-B)** Transient (7h *egr*-expression), and chronic (24h *egr*-expression or 44h *Ras^v12^,scrib-RNAi*-expression) activation of JNK/AP-1 signaling stratifies JNK-activating tissue into distinct JNK/AP-1 and JAK/STAT signaling domains **(A)** and cell behaviors **(B)**. **(C)** *egr*-expressing discs spatially stratify tissue repair behaviors via a mutual repression motif. JNK/AP-1 represses JAK/STAT signaling activitiy via Ptp61F, while JAK/STAT represses JNK/AP-1 activation via Zfh2. JNK signaling ensures G2 arrest and cell survival, while JAK/STAT signaling ensures proliferation and supports survival in the absence of JNK. **(D)** Disruption of the mutual repression motif by cell-autonomous activation of JNK/AP-1 and JAK/STAT signaling overrides the protective G2 stall and drives cells into mitotic cycling and G1, where p53 activation can lead to apoptotic outcomes.

By reducing Stat92E activation upon damage, JNK/AP-1 creates a permissive state for the G2 stall by preventing cycling, thereby protecting cells from apoptosis in a cytotoxic, inflammatory environment (Fig. 7C; Fig. S1A). When high levels of both pathways are co-activated in the same cell, the anti-apoptotic state of the G2 stall is in conflict with proliferative outcomes and therefore active cycling which causes an increase in apoptosis. Consequently, apoptosis of these wound center signaling cells could jeopardize wound organizer function and ultimately tissue repair behaviors (Fig.7D). Therefore, the JNK-activated G2-stall and JAK/STAT-activated cell cycling need to be spatially segregated. Importantly, our observations indicate that, with exceptions [132], even successful tumors avoid coexistence of JNK and JAK/STAT signaling in the same cell as they do not profit from the conflicting cell-autonomous inputs on proliferation and survival decisions. Instead, tumors may highjack a wound organizer network to chronically activate paracrine mitogenic signals but outsource proliferation to other cells as a consequence of the senescent properties of JNK-signaling, G2-stalled cells [31].

We find that spatial patterns generated by the JNK/AP-1 and JAK/STAT regulatory network act at different length scales, depending on the duration and strength of JNK/AP-1 activation (Fig. 7A). Transient or low JNK/AP-1 activity generates short range spatial segregation of JNK/AP-1 and JAK/STAT. Specifically, we observed their segregation between neighboring cells upon transient expression of *egr* or mild disruption of barrier function in *scrib*-RNAi expressing tissue. Cell-by-cell segregation may allow tissue repair processes to respond dynamically. Switching between (1) a stalled, apoptosis-resistant state supporting survival and paracrine signaling in an inflammatory environment or (2) a regenerative, proliferative phenotype could facilitate continuous integration of the repair state. As wound healing progresses and the JNK/AP-1 signaling field shrinks, shifting spatial probabilities of stalled and cycling cells would ultimately resolve the damage.

In contrast, prolonged or high JNK/AP-1 activity segregates JAK/STAT signaling more robustly and consequently into larger bistable fields, prolonging wound resolution and increasing the risk for chronic wound pathologies. Indeed, prolonged JNK/AP-1 activity in *Ras^v12^,scrib*-RNAi tumors or by chronic expression of *egr* drives robust bistable segregation of JNK/AP-1 and JAK/STAT signaling into larger domains. Similarly, chronic wounds in human patients display striking segregation of cellular responses. In a central inflammatory domain, cells acquire senescent phenotypes, arrest and fail to proliferate. In the periphery, a hyperproliferative edge is evidence of chronic mitogenic signals emanating from the wound [3, 9, 10, 133]. These clinical patterns resemble patterning in *egr-*expressing discs, thereby suggesting that *Drosophila* imaginal discs may represent a suitable model to study wound pathologies driven by prolonged activation of inflammatory signaling.

## Materials and Methods

### Fly stocks

All fly stocks and experimental crosses were maintained on standard media and raised at 18 °C or room temperature (22 °C) unless otherwise specified. For detailed genotypes, please refer to Table S1 and S2.

### Fly genetics

Fosmids carrying GFP-tagged *hop* (VDRC 318158) and *Stat92E* (BDSC 38670) alleles, expressed proteins at expected molecular weights and, at least partially complemented respective null alleles, suggesting that both GFP-fusion proteins are functional. A potential Dome-GFP fosmid (VDRC 318098) failed to give rise to a GFP protein by immunofluorescence and Western blot analysis and failed to complement a *dome* allele, thus Dome levels could not directly be analyzed. Genetic complementation assays were carried out by crossing the GFP-tagged fosmids into the background of LOF mutants (*dome^G0441^, hop^9p5^* and *stat^85C9^*). For X-linked *dome^G0441^* and *hop^9p5^* alleles, we calculated the percentage of hemizygous viable adult males emerging from control and experimental crosses (Fig. S4.1A,B). For the autosomal *stat^85C9^* allele, we determined pupariation success of homozygous mutant larvae by counting the number of pupariated larvae in control and experimental crosses (Fig. S4.1C).

We validated RNAi lines by demonstrating that they caused changes to the developmental pattern of JAK/STAT activity in the hinge if expressed throughout normal wing disc development. Crosses were set as previously described, using the *en-GAL4* driver to express the RNAi in the posterior compartment. Larvae were kept at 18°C till D7 AEL. On D7, early L3 staged larvae were dissected, showing characteristic JAK/STAT reporter activity pattern in the hinge. Following dissection and staining, sibling controls and RNAi expressing discs were mounted on the same slide. All discs were imaged at the same settings and the JAK/STAT reporter intensities were quantified in selected regions in the anterior and posterior hinge (Fig. S4.3C). Intensity values from the anterior compartment (not expressing the RNAi) was used as the internal control.

### Flip-out clones

GAL4/UAS-driven ‘flip-out’ overexpression experiments utilized heat-shock-driven expression of a flippase to clonally express *a UAS* construct of choice. After 6h of egg collections, heat-shock was induced on developmental day 5 or 6 at 37°C for 7-10 min. Larvae were dissected at wandering 3rd instar stage or as indicated (28h or 48h after heat-shock, Fig. 2A-D, S2.A-D).

### Genetic cell ablation using Gal4/UAS/Gal80ts

To induce expression of *egr*, experiments were carried out as described in [31, 38, 51] with few modifications. Briefly, larvae of genotype *rn-GAL4, tub-GAL80^ts^* and carrying the desired *UAS-*transgenes were staged with a 6h egg collection and raised at 18°C at a density of 50 larvae/vial. Overexpression of transgenes was induced by shifting the temperature to 30°C for 24h at D7 after egg deposition (AED), and larvae dissected at recovery time point R0 h (Fig. S1.1C) unless noted otherwise. For time course and tumor experiments, transgenes were induced by shifting the temperature to 30°C for 0h, 7h, 14h or 24h at D7 (Fig. S1.2, 2 and S2), or 44h at D6 (Fig. 6 and S6) respectively. Larvae were subsequently dissected for analysis or allowed to recover at 22°C for the indicated time. All images represent R0 h unless noted otherwise. Control genotypes were either *rn^ts^>*, or sibling animals (*+/TM6B, tubGAL80 or +/TM6c*) (Smith-Bolton et al. 2009). All experiments were performed with ≥ 2 biological replicates.

### Immunohistochemistry

Wing discs from third instar larvae were dissected and fixed for 15 min at room temperature in 4% paraformaldehyde in PBS. Washing steps were performed in PBS containing 0.1% TritonX-100 (PBT). Discs were then incubated with primary antibodies (described in **Table S1)** in PBT, gently mixing overnight at 4°C. Tissues were counterstained with DAPI (0.25 ng/µl, Sigma, D9542), Phalloidin-Alexa Fluor 488/647 (1:100, Life Technologies) or Phalloidin-conjugated TRITC (1:400, Sigma) during incubation with cross-absorbed secondary antibodies coupled to Alexa Fluorophores (Invitrogen or Abcam) at room temperature for 2h. Tissues were mounted using SlowFade Gold Antifade (Invitrogen, S36936). Whenever possible, experimental and control discs were processed in the same vial and mounted on the same slides to ensure comparability in staining between different genotypes. Images were acquired using the Leica TCS SP8 Microscope, using the same confocal settings and processed using tools in Fiji. Of note, all wing discs expressing the *10xStat92E>dGFP* reporter were boosted with an anti-GFP antibody.

### EdU Labelling

EdU incorporation was performed using the Click-iT Plus EdU Alexa Fluor 647 Imaging Kit, (described in **Table S1)** prior to primary antibody incubation. Briefly, larval cuticles were inverted in Schneider’s medium and incubated with EdU (10µM final concentration) at RT for 15 minutes. Cuticles were then fixed in 4% PFA/PBS for 15 minutes, washed for 30 minutes in PBT 0.5%. EdU-Click-iT labeling was performed according to manufacturer’s guidelines. Tissues were washed in PBT 0.1%, after which immunostainings, sample processing and imaging were carried out as described above.

### Western Blots

Cell lysates from third instar wing imaginal discs and brains were prepared in lysis buffer (50 mM Tris-HCl pH 7.5, 300mM NaCl, 0.1mM EDTA, 1% Triton X-100, 0.1% SDS, 5% Glycerol, 1mM PMSF, 1/10 tablet of Complete Mini Protease Inhibitor Cocktail) on ice. Protein samples were loaded onto a 10% polyacrylamide gel, along with Chameleon Duo Pre-stained Protein Ladder (LI-COR, P/N 928-60000) as a molecular weight ladder and run at 150V. After SDS-PAGE, proteins were transferred onto a nitrocellulose membrane (Bio-Rad, 162-0115, 0.45µm) in transfer buffer (25 mM Tris, 192 mM glycine, 20% (v/v) methanol) using the wet-tank method with a current density of 300mA for 1h. Prior to antibody incubation, membranes were blocked with 5% milk in PBS. Membrane was incubated with primary antibodies in PBS-T (1% Triton-X in 1xPBS) - rat anti-GFP (Chromotek, 3H9-100, 1:1000) and mouse anti-α-tubulin (Sigma, T9026, 1:5000) on a rotative plate at 4°C overnight. Primary antibodies were removed and washed in dH_2_0 twice for 5 mins each. Membrane was then incubated with secondary antibodies diluted in PBS-T - donkey α-mouse secondary (LI-COR, 926-68072, 1:20000) labelled with a 700nm IRDye and goat α-rat (LI-COR, 926-32219, 1:20000) labelled with an 800nm IRDye. Membrane was washed in dH_2_0 twice for 5 mins each before detection using the Image Studio Software on the Odyssey SA system (LI-COR) (Fig. S4.1D).

### Image Analysis and Quantification

#### General comments

Images were processed, analyzed and quantified using tools in Fiji (ImageJ v2.0.0) [134]. Extreme care was taken to apply consistent methods (i.e. number of projected sections, thresholding methods, processing) for image analysis. Figure panels were assembled using Affinity Designer v1.10. Statistical analyses were performed in Graphpad Prism or R v3.3.3 (www.R-project.org).

#### Total disc area quantification

Maximum intensity projections of z-stacks for each disc were obtained in Fiji. Masks were generated after applying a fixed threshold (10-255) to the DAPI channel and noise was reduced using the ‘Despeckle’ function. Area measurement from the resulting masks were obtained using the ‘Measure’ function.

#### *10xStat92E>dGFP* quantification

Maximum intensity projections of selected confocal sections were taken, excluding the peripodium and carefully chosen to capture signaling activity within the disc proper. In the *egr-*expressing or *egr,transgene*-coexpressing discs, a mask of the pouch domain was generated. Region Of Interest (ROI) outlines were either manually drawn on the DAPI channel to include the pouch and hinge domains till the medial hinge fold or by thresholding Nubbin staining. Centroid co-ordinates were identified using the ‘Measure’ function and marked with a point tool. To measure the fluorescence intensity of the *Stat92E>dGFP* reporter, a square ROI (25×25µm) was placed centrally over the centroid and mean intensity within the ROI was obtained using the ‘Measure’ function in Fiji. (Fig. 4)

#### Dcp-1 area quantification

Maximum intensity projections of stacks for each disc were obtained in Fiji. Masks were obtained by thresholding for signal of interest. Signal to noise ratio was enhanced by ‘Remove outliers’ (bright, radius = 1.5) or ‘Despeckle’ functions. The ‘3D object counter’ function was used to obtain surface area of the generated mask. To control for differences arising from the total size of the discs, ratios between the Dcp-1 area and total WID area were obtained for each disc and reported as percentage values. (Fig. 5C,D,M,N)

#### Tracing JNK, JAK/STAT, EdU and Upd reporters profiles

Partial max projections from confocal stacks of the disc proper were generated, carefully excluding any peripodial signal. For hinge profiles of the JAK/STAT reporter, size-matched discs were chosen and line traces were manually drawn along the hinge using the polygon selection tool. Reporter intensity values from the ‘Plot profile’ function were averaged for 0h, 7h and 14h control and *egr-*expressing discs and plotted. (Fig. S1.2)

For reporter patterns along disc center to disc periphery, JNK/AP-1 masks were made using thresholding tools and ROI outlines were generated. Based on ROIs, the centroid value was determined and *n≥15* tracks were drawn from the centroid outwards towards the hinge for a fixed distance - covering the JNK/AP-1 domain, the JNK/AP-1-JAK/STAT interface, and the JAK/STAT signaling domain respectively. Using the ‘Plot profile’ function in Fiji, the fluorescence intensity values of each reporter was obtained along these tracks and averaged. Reporter patterns for each disc was graphed by scaling the averaged values from each track, between 0 to 1 and plotted on the Y-axis. This was done independently for n≥3 discs, and a representative image was selected for visualization of the spatial trends of the reporters. (Fig 1M,N)

#### Density plots for JNK/AP-1 and JAK/STAT reporters

For combined pouch and hinge regions (Fig. 1I,J), the ROI outlines were manually drawn on the DAPI channel to include the pouch and hinge domains till the medial hinge fold. Partial max projections of z-sections from the disc proper were generated, carefully excluding any peripodial signal.

For time course experiments with *egr-*expressing discs (S2.H-K), a central z-section was chosen and ROIs generated by thresholding the *TRE-RFP* channel to include only the JNK-signaling pouch domain. After applying a ‘Gaussian blur’ filter (sigma=2), masks were generated by applying a low fixed threshold (value = 10) based on early, 7h *egr-*expressing discs. Noise removal was done using the ‘Despeckle’ function, followed by ‘Fill Hole’ function. ROI outlines were generated using ‘Create selection’. All ROIs were placed on the source *TRE>RFP* and *Stat92E>dGFP* reporter channels. ‘Save X-Y co-ordinates’ function was used to obtain pixel fluorescence intensity values for both reporters within the ROI, and the distribution of fluorescence intensities across the samples were graphed by generating a binned (bins = 16) 2D histogram plot using R. After all NA values were set to 0, data density per bin for each disc was calculated as a percentage. Values were averaged across corresponding bins for all discs within a timepoint to generate the average fluorescence intensity distribution profile. To distinguish regions with low, medium and high signaling for each pathway, the graphs were subdivided into sections s1-s9 by defining a threshold to distinguish low, medium and high JNK/AP-1 and JAK/STAT signal intensities (Fig. S2K). Percentage of datapoints within each section was determined from the averaged fluorescence intensity distribution profile. Ordinary 2-way ANOVA with Dunnett’s multiple comparisons test was carried out for statistical analysis. Alpha was set at 0.05

#### Cell cycle profiles

Representative sections were selected per disc and a square ROI area (75×75µm^2^) was placed over the central pouch cells. Only viable cells stained with characteristic euchromatic and heterochromatic DAPI staining were chosen. Using the ‘Multi point’ tool, cells showing a G1 profile (green), G2 profile (yellow+red) or neither (black) were counted. Percentage values for the gap phases were calculated for n≥3 discs per genotype and visualized. T-tests were performed on the calculated G1:G2 ratios per disc. (Fig 5G,H)

#### JAK/STAT reporter quantifications for testing RNAi lines

Crosses were setup as previously described and wing discs from larvae were dissected on D7 AED. Single sections at similar focal planes were carefully chosen during imaging. Using drawing tools in Fiji, ROIs were generated to include JAK/STAT signaling cells in the hinge within A and P compartments. The mean JAK/STAT reporter intensity within the ROI was obtained from control and RNAi expressing discs using the ‘Measure’ function in Fiji. For each dataset, paired t-tests were performed between A and P fluorescence intensity values for the same genotype. (Fig. S4.3A-C)

#### Nuclear translocation of Stat92E-GFP

Images from the pouch (n=12) and hinge (n=9) domains of control discs and pouch (n=16) and hinge (n=12) domains of *p35+egr* expressing discs were obtained to track the intra-cellular localization of *Stat92E-GFP* within the nucleus and cytoplasm. Images were taken at high magnification (63x). After thresholding (‘Huang’), the selection tool was used to generate a nuclear ROI using the DAPI channel. The inverse selection tool was used to generate a corresponding cytoplasmic ROI for each image (Fig S4.1L). These nuclear and cytoplamic ROIs were then placed on the *Stat92E-GFP* channel and mean fluorescence intensity values were obtained for each subcellular fraction and unpaired t-tests were performed. (Fig. S4.1M)

#### MMP-1 intensity quantification

To measure the fluorescence intensity of MMP-1 in *egr-*expressing and *egr,zfh2-co*expressing discs, maximum intensity projections of confocal sections were taken. Sections were carefully chosen to capture JNK/AP-1 signaling activity in the disc proper, avoiding the peripodium and cell debris. Square ROIs of fixed length (30 µm) were placed in 3 non-overlapping regions in the pouch domain, and mean intensity was calculated using the ‘Measure’ function in Fiji. Mean fluorescence intensities obtained from ROIs were averaged per disc and values from each disc per genotype were used for statistical analysis. (Fig. S3A-C)

#### JNK^+^ and JNK^-^ Area and intensity quantifications (*Ras^v12^,scrib-RNAi* tumors)

A representative z-section was selected per disc. As our interest was in quantitating the spatial segregation, only JNK/AP-1 or JAK/STAT signaling regions in the tumor were chosen. After applying a ‘Gaussian blur’ filter (sigma=2), ‘Moments’ thresholding and ‘Despeckle’ function for noise correction was applied to generate masks for regions showing *TRE>RFP* (JNK) and *Stat92E>dGFP* (JAK/STAT) reporter activity. From the ‘Image calculator’ function, ‘AND’ and ‘Subtract’ operations on JNK/AP-1 and STAT masks were carried out to obtain the JNK^+^STAT^+^ mask, and the exclusive JNK^+^ (JNK-STAT) or exclusive JNK^-^(STAT-JNK) masks respectively. Area of each mask was obtained using the ‘Measure’ function. Summed area of these masks were considered as the total area and each area was then represented as a percentage of total. One-way paired ANOVA with Holm-Šídák’s multiple comparison test was performed to test for statistical significance. Mean intensity for *TRE>RFP* or *10xStat92E>dGFP* reporters were measured inside the JNK/AP-1 mask (JNK^+^) and the exclusive JNK^-^ masks. Paired t-tests were performed to test for statistical significance. (Fig. 6E,F and S6I)

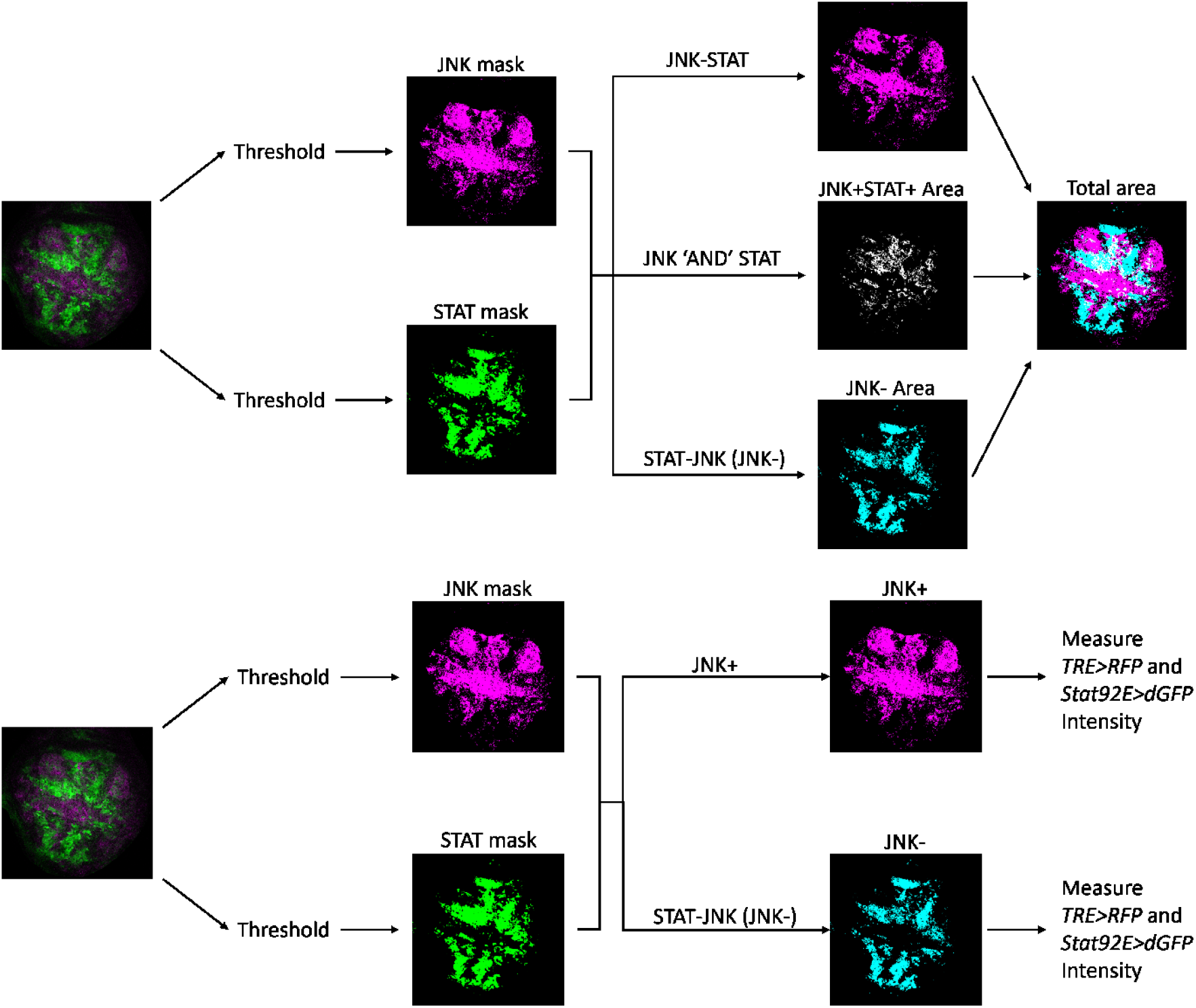

#### ph3 counts

Partial maximum intensity projection of three central z-sections were taken per disc. To generate masks for regions showing *Stat92E>dGFP* reporter activity (STAT^+^), a ‘Gaussian blur’ filter (sigma=1), ‘Moments’ threshold and ‘Despeckle’ function for noise correction was applied. The DAPI channel was used to manually outline the pouch domain and generate masks. Using the ‘Image calculator’ function, STAT^+^ mask was ‘Subtract’-ed from the pouch mask to generate the STAT^-^ mask. Areas of the STAT^+^ and STAT^-^ regions in the pouch were obtained using the ‘Measure’ function. Then, the ph3 channel was duplicated and ‘Li’ threshold was applied. ‘Remove outlier’ (radius=5 threshold=50) and ‘Despeckle’ function for noise correction, followed by ‘Watershed’ function was applied to generate a ph3 mask. Using the ‘AND’ operator, the ph3 mask was overlayed with the STAT^+^ and STAT^-^ masks. ‘3D object counter’ function was then used to obtain ph3 ‘Object counts’ within the STAT^+^ and STAT^-^ regions and divided by area of each ROI to get the ph3 counts per mm^2^. Paired t-tests were performed to test for statistical significance. (Fig. 6I)

### Mathematical modelling

To test for the existence of a regulatory motif within our signaling network, a mathematical model was derived to describe the temporal dynamics of the concentration of specific molecules over a fixed 2D space. The system was modeled as a set of ordinary differential equations (ODEs) (see also Fig. 3C). Two sets of modeling equations were established, that reflected conditions which include or exclude repression of JAK/STAT by JNK. The term describing repression of JAK/STAT (referred to as ‘JAK’ in the modeling equations and hereafter) by JNK/AP-1 is highlighted in red.

Uni-directional repression model (see also Fig. 3B):

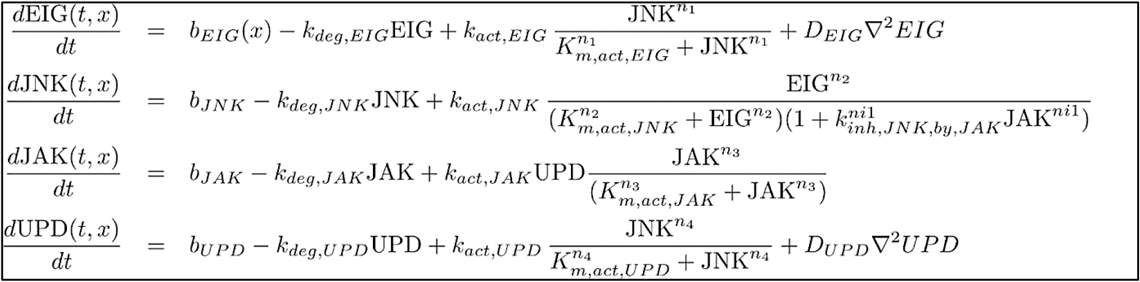

Mutual repression model (see also Fig. 3B’):

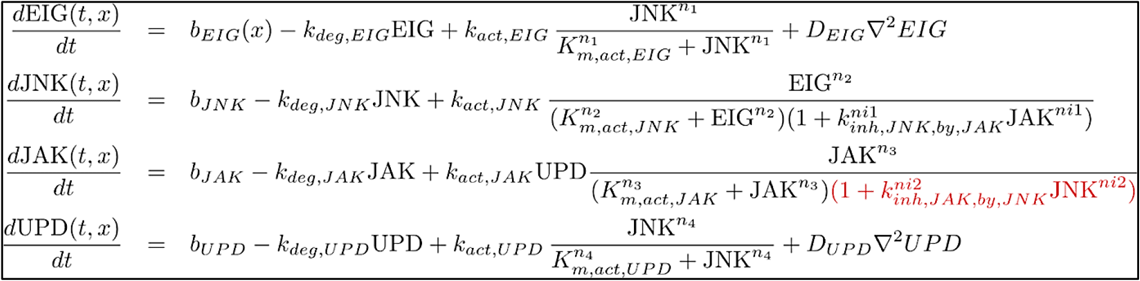

Characteristic units are represented by state (u), space (x) and time (t). The mathematical terms include, among others, basal rate of production (*b*), activation rate (*k_act_*), linear degradation rate (*k_deg_*), Michaelis–Menten constant (*K_m_*), inhibition constant (*k_inh_*), diffusion co-efficients (*D_EIG_, D_UPD_*), Hill co-efficients to model cooperativity (*n_1_,n_2_,n_3_,n_4,_ ni_1_,ni_2_*) and Hill kinetics to describe self-amplification and mutual repression.

The first equation describes the dynamics of the concentration of Egr, produced at a basal rate dependent on the spatial co-ordinate x and degraded at a linear rate. Hill kinetics describe the production of Egr induced by JNK/AP-1 and its diffusion away from the source. The second equation describes the dynamics of the concentration of the JNK/AP-1 TF, produced at a basal rate and degraded linearly. Hill kinetics describe production of JNK/AP-1 induced by *egr* as well as inhibition of JNK/AP-1 by JAK [51]. The third equation describes the dynamics of the concentration of the JAK TF, produced at a basal rate and degraded linearly. Hill kinetics describe positive feedback dependent on Upd. Inhibition of JAK by JNK/AP-1 is described and the model is tested with and without this mathematical term (highlighted in red) to check if this interaction exists within the system. The fourth equation describes the dynamics of the concentration of Upd, produced at a basal rate and degraded linearly. Hill kinetics describe the production of Upd induced by JNK/AP-1 and its diffusion [28, 48, 51].

Technically, the modeling was done based on a non-dimensionalized version of the model, rescaled to reduce the number of free parameters. The system was then defined as follows:

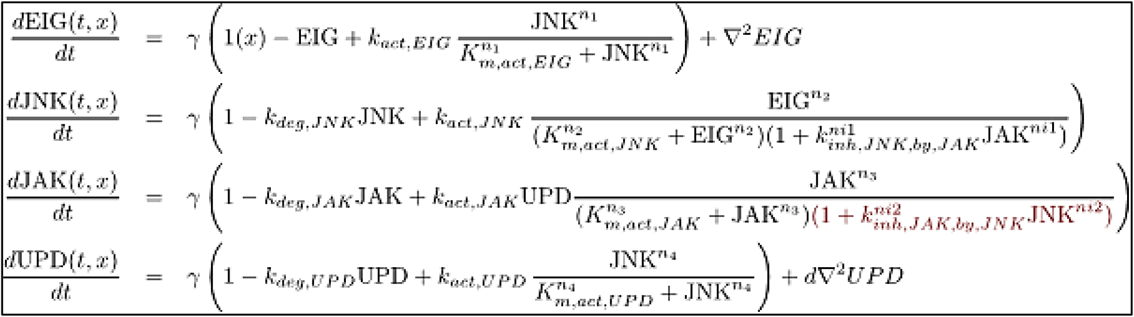

For each dimensionless parameter, the parameter dependencies and the scanned parameter range are represented in the table below:

| Dimensionless parameter | Functional relation to systems parameter | Scanned parameter range |
| --- | --- | --- |
| $\gamma$ | $k_{deg, EIG} L^2 / D_{EIG}$ | 0.01 ... 10000 |
| $\hat{k}_{deg, JNK}$ | $k_{deg, JNK} / k_{deg, EIG}$ | 0.01 ... 10000 |
| $\hat{k}_{deg, JAK}$ | $k_{deg, JAK} / k_{deg, EIG}$ | 0.01 ... 10000 |
| $\hat{k}_{deg, Upd}$ | $k_{deg, Upd} / k_{deg, EIG}$ | 0.01 ... 10000 |
| $\hat{k}_{act, EIG}$ | $k_{act, EIG} / b_{EIG}$ | 0.00001 ... 10 |
| $\hat{k}_{act, JNK}$ | $k_{act, JNK} / b_{JNK}$ | 0.01 ... 10000 |
| $\hat{k}_{act, JAK}$ | $k_{act, JAK} D_{EIG} / L^2 / b_{UPD} / b_{JAK}$ | 0.01 ... 10000 |
| $\hat{k}_{act, UPD}$ | $k_{act, UPD} / b_{UPD}$ | 0.01 ... 10000 |
| $\hat{K}_{m, act, EIG}$ | $K_{m, act, EIG} D_{EIG} / L^2 / b_{EIG}$ | 0.00001 ... 10 |
| $\hat{K}_{m, act, JNK}$ | $K_{m, act, JNK} D_{EIG} / L^2 / b_{JNK}$ | 0.01 ... 10000 |
| $\hat{K}_{m, act, JAK}$ | $K_{m, act, JAK} D_{EIG} / L^2 / b_{JAK}$ | 0.01 ... 10000 |
| $\hat{K}_{m, act, UPD}$ | $K_{m, act, UPD} D_{EIG} / L^2 / b_{UPD}$ | 0.01 ... 10000 |
| $\hat{k}_{inh, JAK, by, JNK}$ | $k_{inh, JAK, by, JNK} b_{JAK} L^2 / D_{EIG}$ | 0.01 ... 10000 |
| $\hat{k}_{inh, JNK, by, JAK}$ | $k_{inh, JNK, by, JAK} b_{JNK} L^2 / D_{EIG}$ | 0.01 ... 10000 |
| $\hat{n}_i$ | $n_i$ | 1 ... 4 |
| $d$ | $D_{UPD} / D_{EIG}$ | 1.5 ... 4 |

We performed a global analysis by sampling approach introduced in *Rausenberger et. al* [135]. The models were simulated on a 1D spatial domain x = [0 1]. To represent the wound site, a high basal production of Egr was fixed within a finite space at x = [0 0.05], while basal rate of production of Egr for the rest of the region was set to zero. The rate of production of Upd was defined as a function of JNK/AP-1 activity [18, 20, 39, 51]. For all states, initial condition was set as u (t=0,x) = 0.1. Solutions for each model (with and without repression) were simulated using 10^6^ parameter sets, drawn uniformly from an interval from 10^-3^ to 10^+2^ on a logarithmic scale, and tested under different conditions of diffusion – *D_Upd_ > D_Egr_, D_Upd_ < D_Egr_* and *D_Upd_ = D_Egr_*. The resulting solutions were automatically scored for countergradient endpoint features extracted from the experimentally observed gradient patterns.

Using this approach [135], positive solutions for a generic bistable pattern from the scanned parameter range were automatically extracted based on the following features:

i. JAK at x = 0 lower than JAK at x = 1
ii. JNK/AP-1 at x = 0 higher than JNK/AP-1 at x = 1
iii. Relative difference of JAK higher than 10%
iv. Relative difference of JNK/AP-1 higher than 10%

Positive solutions for a specific bistable pattern which mimics biologically observed patterns were automatically extracted based on the additional features:

v. JAK has a local maximum which is not at the boundary of the simulated domain
vi. Height of local JAK maximum at least 5%

## Author Contributions

Conceptualization JJ, VW, AKC

Validation JJ, AA, IG, AKC

Investigation JJ, RE, AA, CC, AKC

Writing JJ, VW, RE, JT, AKC

Visualization JJ, RE, VW, AKC

Supervision JT, AKC

Funding Acquisition AKC

## Acknowledgements

We thank the LIC facility (University of Freiburg) for technical help with imaging. We thank Erika Bach, David Bilder, Dirk Bohmann, Stephen Cohen, Fernando Diaz-Benjumea, Iswar Hariharan, Norbert Perrimon, Helena Richardson, Reinhardt Schuh, Y. Henry Sun, Mirka Uhlirova and Pelin Volkan for sharing reagents and the Bloomington Drosophila Stock Center (BDSC), the Vienna Drosophila Stock Collection (VDRC), the University of Zurich ORFeome Project (FlyORF), the Developmental Studies Hybridoma Bank (DSHB) and the Monoclonal Antibody Core Facility at the Helmholtz Zentrum Munich for providing fly stocks and antibodies. Thanks to Florian Steinberg (University of Freiburg) for support with protocol, reagents and imaging equipment for Western blot assays. We also thank the SGBM graduate school for support.

---

**Figure S1.1.**
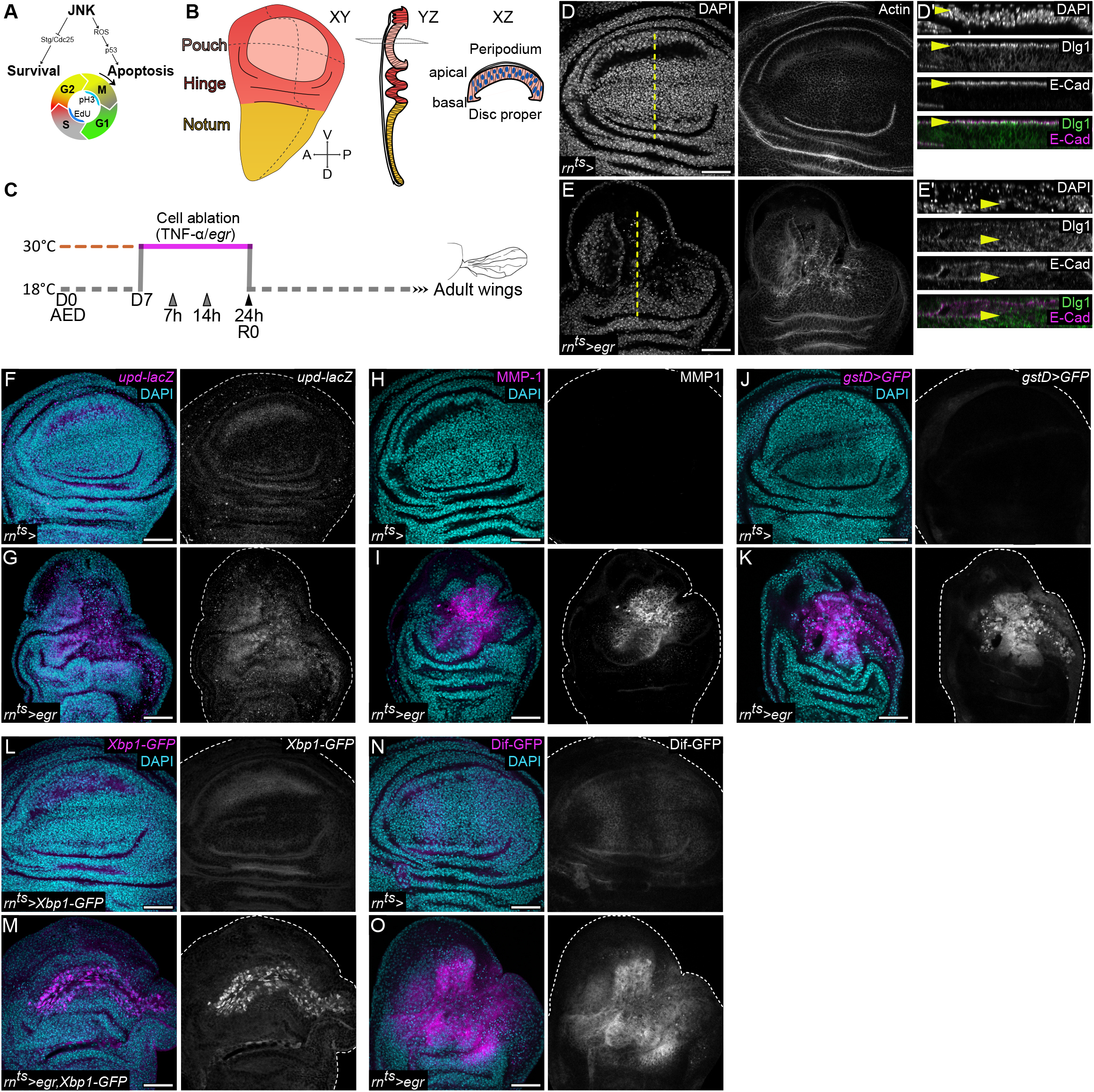
*egr*-expressing wing discs display hallmarks of tissue inflammation. **(A)** Schematic of the FUCCI toolkit, utilizing the degradable *mRFP-NLS-CycB*^1-266^ (red) *and GFP-E2F1*^1-230^ (green) reporters as readouts for G1 and late S/early G2 phases of the cell-cycle respectively. Cells with elevated levels of both FUCCI reporters (yellow) are in the late G2 phase of the cell-cycle (Zielke et al. 2014). Cell in the S-phase and M-phase are detected using EdU and PH3 respectively. Damage-induced JNK/AP-1 signaling mediates survival or apoptosis through control of control of cell cycle progression (Cosolo et al. 2019). **(B)** Schematic of a central XY, YZ and XZ section through a third instar wing imaginal disc. Different shades in the XY section represent the pouch, hinge and notum domains. Dashed lines represent the anterior-posterior (A/P) and dorsal-ventral (D/V) compartment boundaries. The YZ section visualizes the pseudostratified monolayer organization of the disc proper, and the overlying peripodial epithelium layer. XZ section visualizes the apical to basal orientation of epithelial cells in the wing pouch. **(C)** Experimental timeline denoting the protocol used for induction of cell-ablation in this study. Larvae were raised at 18°C, and on D7 After Egg Deposition (AED), vials were transferred to 30°C to induce expression of pro-apoptotic transgenes for 24 h in the wing imaginal discs (also referred to as Recovery timepoint 0 or R0), using the pouch-specific *rotund-GAL4* (*rn-GAL4*) driver. Gray arrowheads represent additional recovery points used for time-course experiments. Shifting the larvae back to 18°C terminates expression of pro-apoptotic transgenes and thus ablation. The prospective adult wing blade arising from the damaged wing pouch allows for assessment of regenerative capacity. **(D-E’)** XY view of control **(D)** and *egr-*expressing **(E)** discs at R0 stained for actin. XZ-sections through the tissue **(D’,E’)** were visualized along dotted yellow lines. Immunostaining for E-cadherin (E-Cad, magenta) and Discs large (Dlg, green) reveals loss of barrier integrity (yellow arrowheads) in an *egr*-expressing disc **(E’)** when compared to a control disc **(D’)**. **(F-O)** Control **(F,H,J,L,N)** and *egr*-expressing **(G,I,K,M,O)** discs at R0. *egr-*induced JNK/AP-1 signaling results in strong upregulation of various markers associated with inflammation and senescent properties – cytokines – *upd*-LacZ **(F,G)**, metalloproteases – MMP-1 **(H,I)**, ROS response – *gstD*>GFP **(J,K)**, UPR – spliced Xbp1-GFP **(L,M)** and NF-kB signaling – Dif-GFP **(N,O)**. Maximum projections of multiple confocal sections are shown in **F-O**. Discs were stained with DAPI to visualize nuclei. Scale bars: 50 µm

**Figure S1.2.**
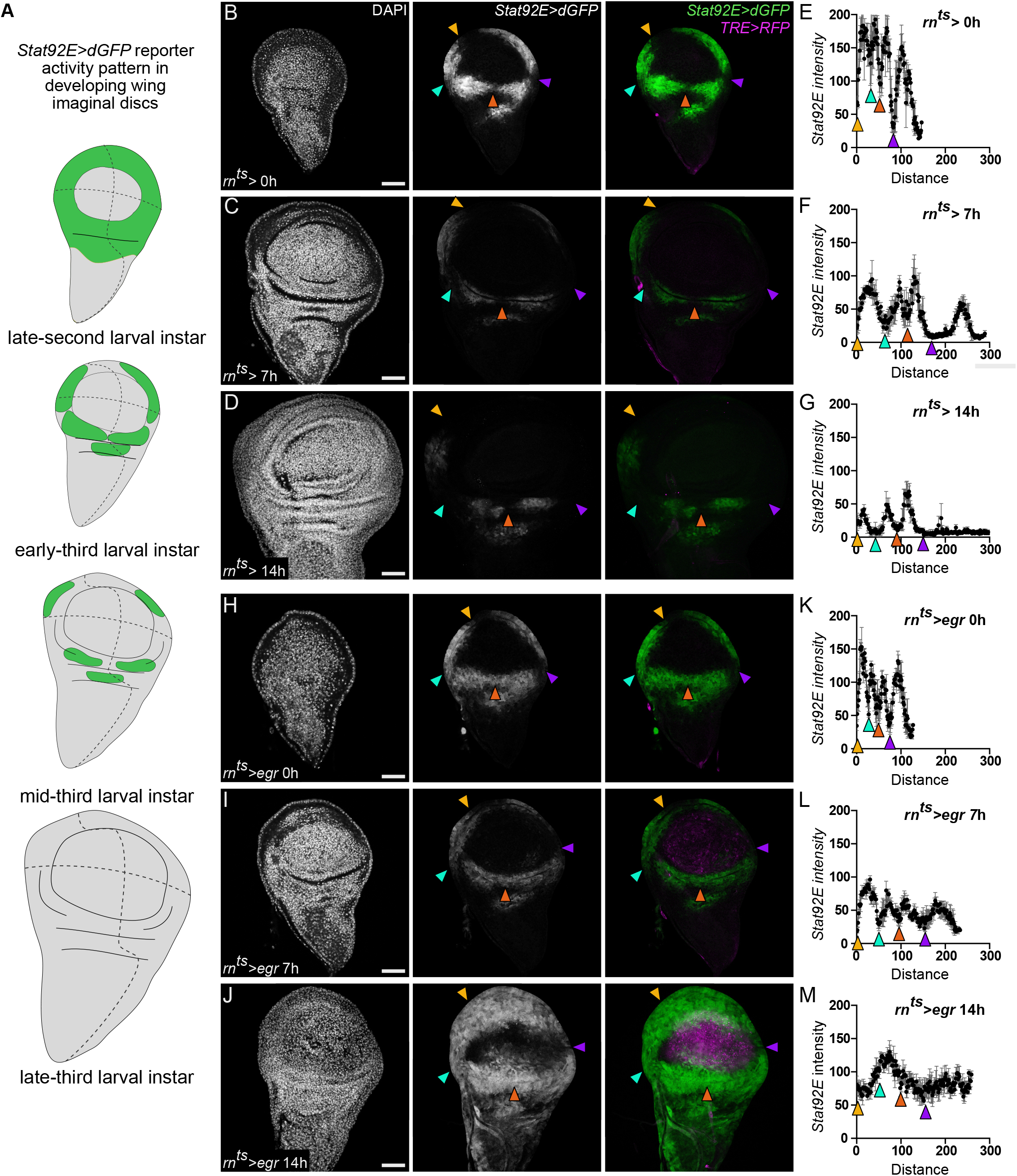
JAK/STAT is activated *de novo* downstream of JNK/AP-1 signaling upon tissue damage. **(A)** Schematic showing dynamic patterning of the JAK/STAT signaling reporter (*Stat92E>dGFP,* green) activity in developing, undamaged wing imaginal discs. Note the gradual loss of hinge-specific patterns from mid to late third larval instar stages. **(B-D)** Undamaged control discs at 0h **(B)**, 7h **(C)** and 14h **(D)** visualized for JNK/AP-1 and JAK/STAT reporter activity patterns using the *TRE>RFP* (magenta) and *Stat92E>dGFP* (green) reporters. **(H-J)** Wing discs after 0h **(H)**, 7h **(I)** and 14h **(J)** of *egr*-expression in the pouch domain. Increasing JNK/AP-1 reporter activity is seen in the pouch from 7h to 14h of *egr*-expression. **(E-G, K-M)** *Stat92E>dGFP* reporter fluorescence intensity, traced along the hinge of control and *egr*-expressing discs at 0h **(E,K)**, 7h **(F,L)** and 14h **(G,M)**. Colored arrowheads indicate similar positions along hinge regions where JAK/STAT reporter activity is progressively lost. Note the *de novo* increase in the mean JAK/STAT reporter intensity in the hinge domain of *egr-*expressing discs **(M)** compared to control, undamaged discs **(G)** at 14h. Graphs display mean ± SEM for control 0h, *n=2*; 7h, *n=2* and 14h, *n=2* discs and *egr* 0h, *n=2*; 7h, *n=4* and 14h, *n=5* discs. Maximum projections of multiple confocal sections are shown in **B-D**, **H-J.** Discs were stained with DAPI to visualize nuclei. Scale bars: 50 µm

**Figure S2.**
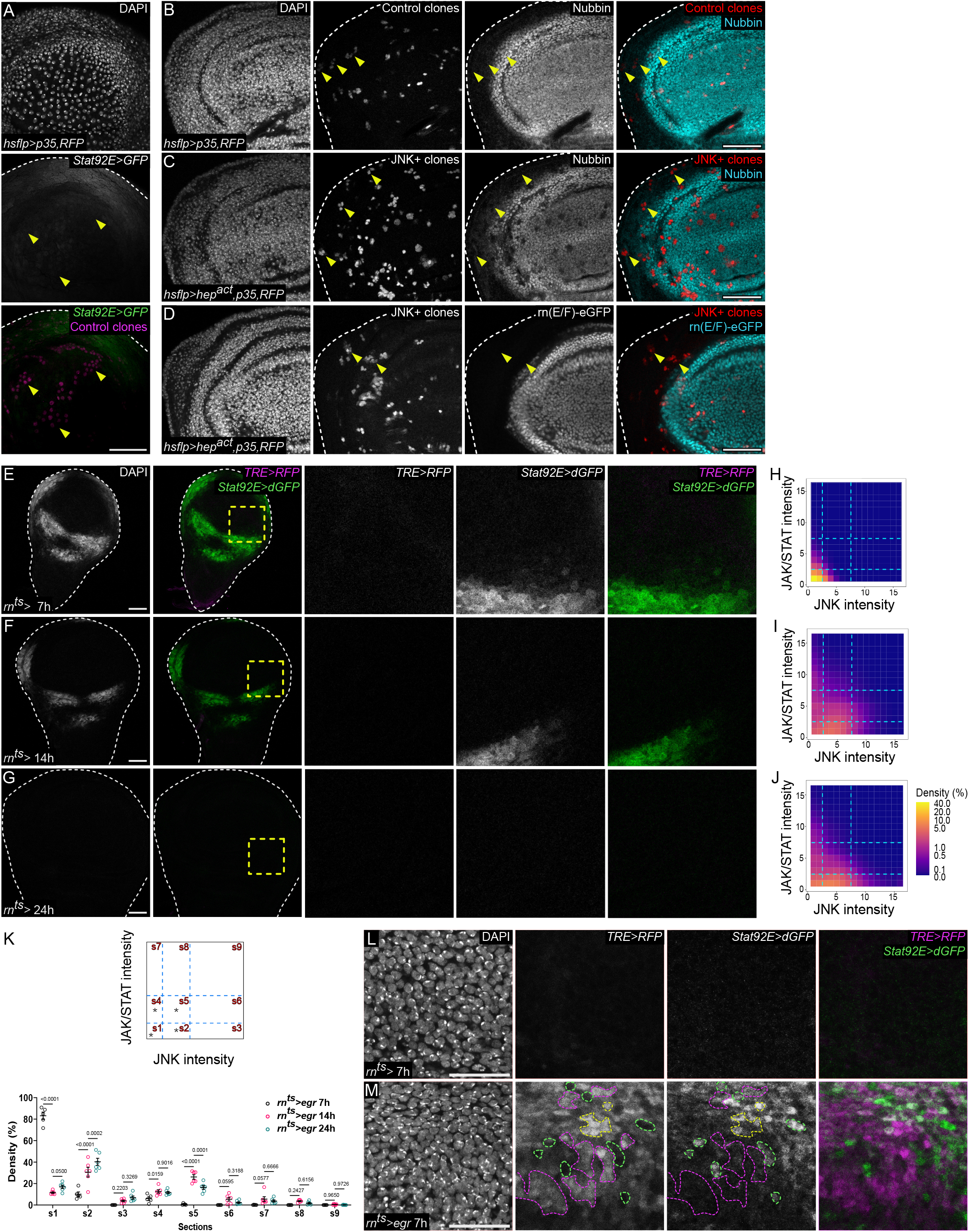
JNK/AP-1 represses JAK/STAT activity independent of pouch-specific transcription factors. **(A)** RFP expression marks *p35*-expressing control clones in the peripodium (magenta) 28h after clone induction in discs expressing the *Stat92E>GFP* reporter (green). Yellow arrowheads indicate clones of interest. Note the absence of ectopic/non-cell-autonomous *Stat92E>GFP* reporter activity around the control clones **(A)** when compared *p35,hep^act^*-expressing JNK+ clones in Figure 2C. **(B-C)** *p35*-expressing control clones (red, **B**) and *p35,hep^act^*-expressing JNK+ clones (red, **C**) 28h after clone induction, stained for the pouch-specific TF Nubbin (cyan). JNK/AP-1 signaling clones show no ectopic expression of Nubbin in hinge, however, Nubbin is completely repressed intra-clonally within the pouch. **(D)** *p35,hep^act^*-expressing clones (red) in a disc expressing the Rn(E/F)-eGFP reporter (cyan) as a readout for the expression of the pouch-specific TF Rotund (Rn) 28h after clone induction. JNK/AP-1 signaling clones show no ectopic intra-clonal expression of Rn in the hinge domain. Yellow arrowheads indicate clones in the hinge. **(E-G)** A time-course analysis of *TRE>RFP* (magenta) and *Stat92E>dGFP* (green) reporter activity in *rn*-expressing control discs after 7h **(E)**, 14h **(F)** and 24h **(G)** of inductive temperature shift to 30°C. Yellow squares show magnified regions. **(H-J)** 2D density plots of averaged pixel fluorescence intensity for the *TRE>RFP* (JNK intensity,) and *Stat92E>dGFP* (JAK/STAT intensity) measured exclusively within the JNK/AP-1 signaling domain (magenta region in Figure 1M) of 7h **(H)**, 14h **(I)** and 24h **(J)** *egr*-expressing discs. Data was binned with a bin size of 16 and X and Y-axis denote the number of bins. Dashed lines (cyan) represent visually defined thresholds for distinguishing low, intermediate and high reporter activity regions. Plots represent averaged density values per bin across 7h, *n=5*; 14h, *n=6* and 24h, *n=6* discs. The same density color scale applies to all genotypes. **(K)** Schematic showing different sections (s1-s9) from which mean pixel density values were obtained for 7h, 14h and 24h discs for statistical analysis. Black asterisks mark sections of interest. Graph comparing mean density of pixels in each section (s1-s9) for different durations of cell ablation Ordinary 2-way ANOVA with Dunnett’s multiple comparisons test was used to test for statistical significance. Graphs represent mean ± SEM for 7h, *n=5*; 14h, *n=6* and 24h, *n=6* discs. Note the reduction in medium JAK/STAT activity within JNK/AP-1 signaling cells in S5 from 14h (15.37%) to 24h (5.25%) in *egr*-expressing discs. **(L-M)** Selected regions from a control wing disc pouch **(L)** and a pouch after 7h of *egr-*expression **(M)** expressing the *TRE>RFP* (magenta) and *Stat92E>dGFP* (green) reporter. Dashed outlines (magenta) of cells activating JNK/AP-1 show an absence of JAK/STAT activity. Dashed outlines (green) cells activating JAK/STAT show an absence of the JNK/AP-1 activity. Dashed outlines (yellow) show a small proportion of cells that can strongly activate both the JNK/AP-1 and JAK/STAT reporters cell-autonomously. Maximum projections of multiple confocal sections are shown in **B-D, L-M** Discs were stained with DAPI to visualize nuclei. Scale bars: 50 µm

**Figure S3.**
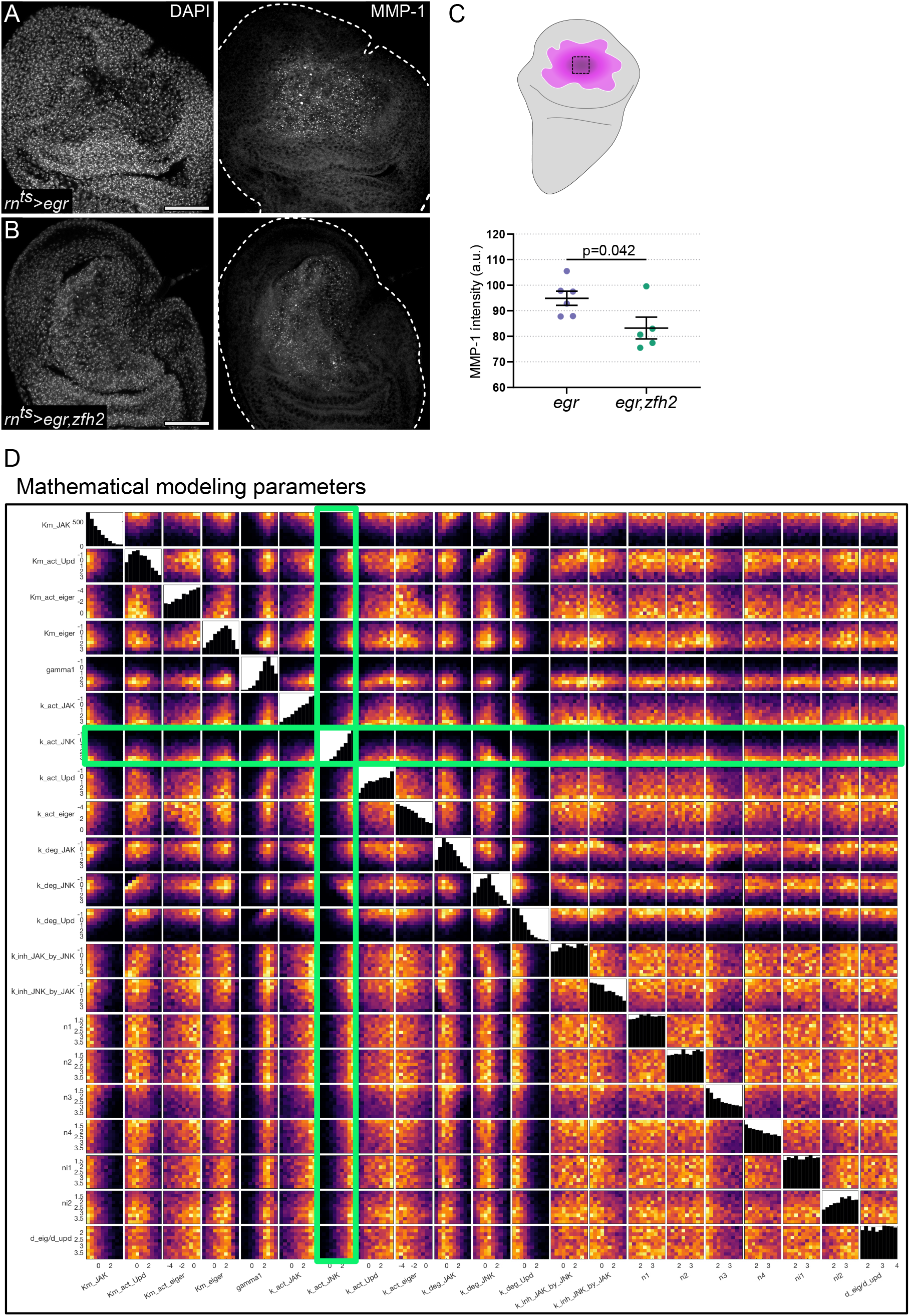
A mutual-repression loop promotes bistable segregation of JNK/AP-1 and JAK/STAT activation patterns. **(A-C)** An *egr*-expressing **(A)** and *egr,zfh2*-co-expressing disc **(B)** at R0, stained for MMP-1. Overexpression of Zfh2 leads to decreased JNK/AP-1 signaling in the central pouch region, as assayed by MMP-1 staining. **(C)** Quantification of MMP-1 fluorescence intensity measured within the JNK/AP-1 signaling pouch domain. Black square in schematic shows measured region. Graphs display mean ± SEM for *n=6*, *egr*-expressing and *n=5*, *egr,zfh2*-co-expressing discs. T-tests were performed to test for statistical significance. **(D)** 21 parameters, their numerical distributions and influence on positive bistable solutions obtained from the mutual repression model (Figure **3B’**) where JNK/AP-1 represses JAK/STAT. Parameters represent reaction rates - for activation, degradation and inhibition of simulated components, as well as hill coefficients and diffusion coefficients for different variables of the model. Bar graphs show the frequency of positive solutions within the tested range of values for each parameter. Our model supports that higher activation rates of JNK/AP-1 (kact JNK) have a strong influence on the total number of positive bistable solutions, highlighted in green rectangles. Maximum projections of multiple confocal sections are shown in **A-B** Discs were stained with DAPI to visualize nuclei. Scale bars: 50 µm

**Figure S4.1.**
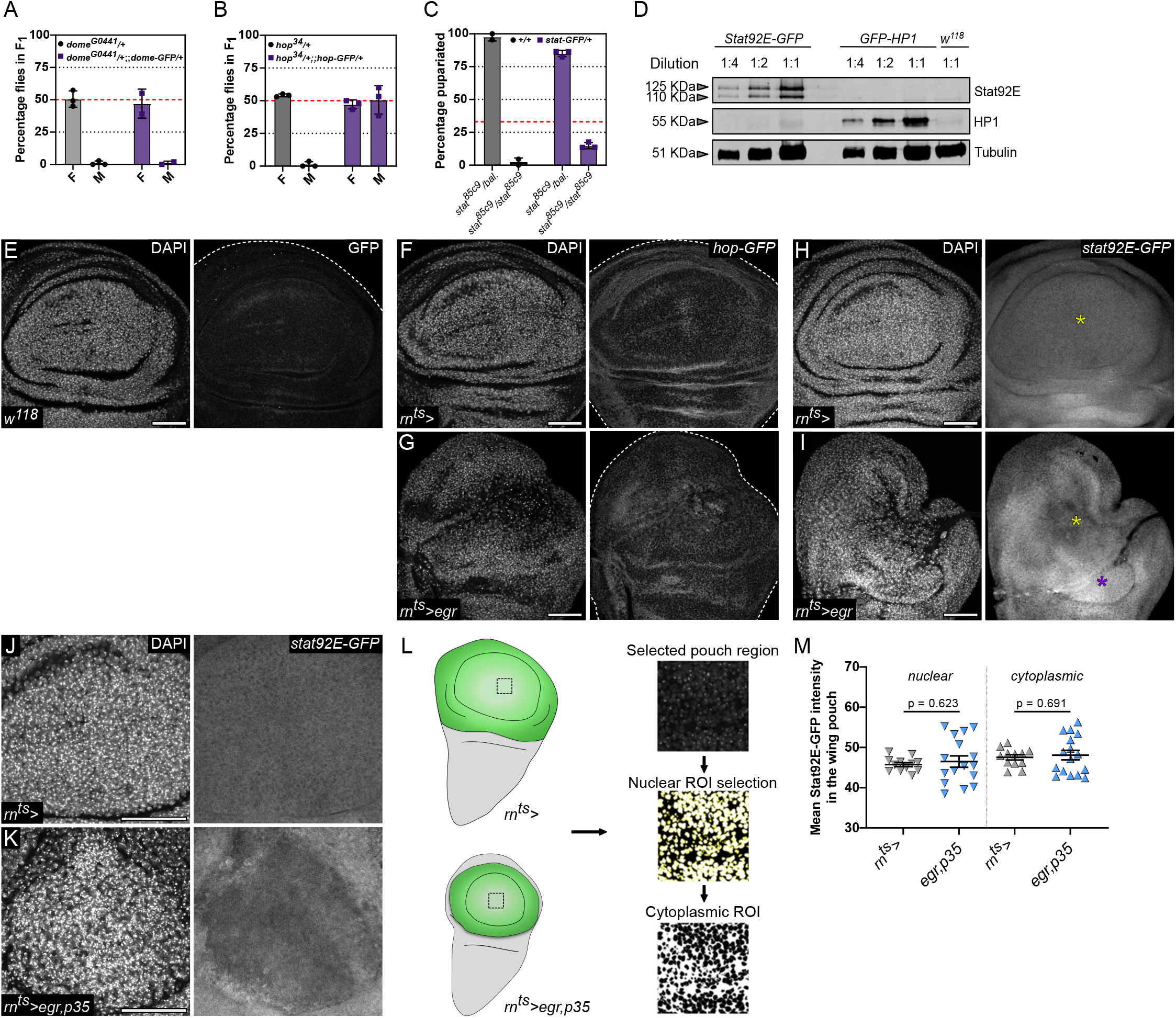
Core components of the JAK/STAT pathway are not altered in JNK/AP-1 signaling cells. **(A-C)** Test for genetic complementation of the LOF mutants *dome^G0441^, hop^9p5^* and *stat^85C9^* (grey bars) by GFP-tagged Dome, Hop and Stat92E fosmid lines (purple bars) (see Experimental procedures). Dome-GFP is unable to rescue male lethality in *dome^G0441^* LOF background to the expected rate of rescue, i.e. 50% of adults in F1 (red dashed line), *n=2* independent replicates **(A)**. Hop-GFP successfully rescues male lethality in *hop^9p5^* LOF background from 0% to 47% of viable adults, at the expected rate of rescue in F1 (red dashed line), *n=3* independent replicates **(B)**. Stat92E-GFP partially rescues larval lethality in *stat^85c9^* null mutants from 0% to 15% pupariation, at approximately half the expected rate of a full rescue (red dashed line), *n=3* independent replicates **(C)**. Thus, the Hop-GFP and Stat92E-GFP fosmid lines were used as proxies for further analysis of protein localization in *egr*-expressing discs. **(D)** Western blots analyzed for Stat92E-GFP expression in imaginal discs. HP-1-GFP genotypes were included as positive control, along with a non-tagged negative control from wild type imaginal disc extracts. Increasing concentrations were loaded, and probed with anti-GFP and anti-Tubulin antibody. The GFP-tagged Stat92E protein is detected at the expected MW (125Kda and 110Kda), and likely representing overlapping isoforms running in two separate weight ranges (71.2-76.8 kD and 85.6-92.8 kD plus GFP-tag). **(E)** A control *w^118^* disc used as a negative control to determine anti-GFP antibody background. **(F-G)** A control **(F)** and *egr-*expressing disc **(G)** expressing the GFP-tagged Hop protein. **(H-I)** A Control **(H)** and *egr*-expressing disc **(I)** expressing the GFP-tagged Stat92E protein. While Stat92E expression is low in the medial hinge folds of control discs, the hinge shows elevated expression in *egr* discs (purple asterisk). This is consistent with JAK/STAT activity inducing expression of Stat92E to promote self-activation. In contrast, Stat92E-GFP protein levels remains low in the pouch (yellow asterisk). **(J-K)** A control pouch **(J)** and a larger domain of *p35,egr-*expressing cells in the pouch **(K)** was used to test for changes in nuclear versus cytoplasmic localization of Stat92E-GFP within high JNK/AP-1-signaling cells. **(L)** Schematic showing the centrally selected region (black square) in control and *egr,p35*-expressing discs, used to measure Stat92E-GFP fusion protein intensity in the JNK/AP-1 signaling pouch domain. Automated workflows were used to generate nuclear and cytoplasmic ROIs from selected regions, to quantify Stat92E-GFP protein levels within these area fractions. **(M)** Stat92E-GFP reporter fluorescence intensity within nuclear and cytoplasmic area fractions of the JNK/AP-1 signaling pouch, calculated for *n=12* independent regions from control and *n=16* independent regions from *egr,p35*-expressing discs. The pouch signaling domain in *egr,p35*-expressing discs fails to exhibit an increase in nuclear translocation of the Stat92E-GFP protein, indicative of basal levels of JAK/STAT signaling in these cells similar to controls. The surrounding hinge domain recapitulated elevated JAK/STAT signaling by elevated Stat92E-GFP protein expression and nuclear translocation (data not shown). Maximum projections of multiple confocal sections are shown in **H-K** Discs were stained with DAPI to visualize nuclei. Scale bars: 50 µm

**Figure S4.2.**
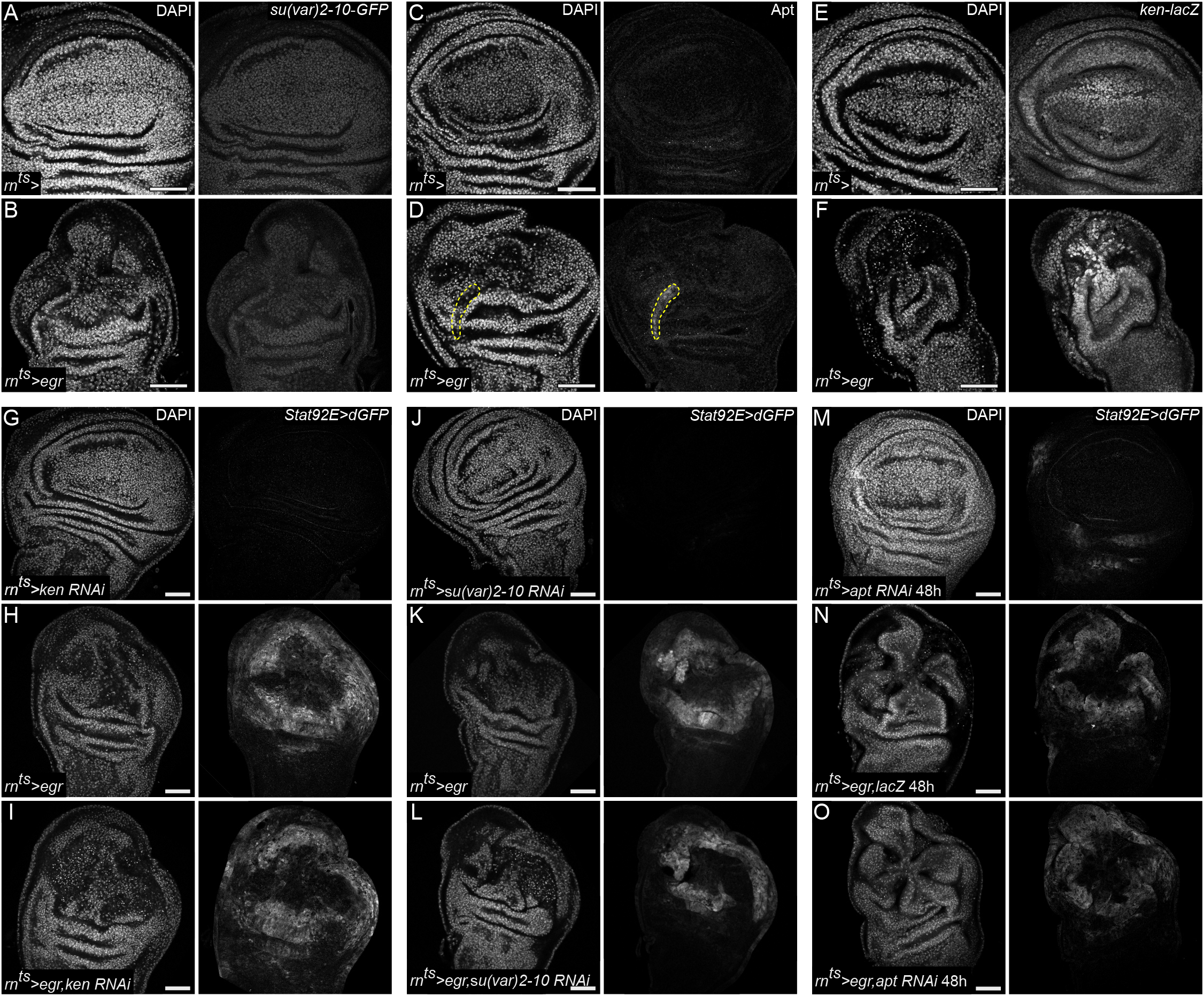
Known negative regulators of JAK/STAT signaling are not required for JNK/AP-1-mediated repression of JAK/STAT. **(A-B)** A control **(A)** and *egr*-expressing disc **(B)**, also expressing GFP-tagged Su(var)2-10/dPIAS and stained for GFP. **(C-D)** A control **(C)** and *egr*-expressing disc **(D)** stained for Apontic (Apt). Yellow outlines mark non-specific tracheal staining in **D**. **(E-F)** A control **(E)** and *egr*-expressing disc **(F),** also expressing *ken*-*Lac*Z and stained for anti-β-Galactosidase. **(G-O)** *RNAi*-expressing control discs **(G,J,M)**, *egr*-expressing **(H,K,N)** and *egr,RNAi*-co-expressing discs **(I,L,O)** visualized for *Stat92E>dGFP* reporter activity. *egr, ken RNAi* **(I)** and *egr, su(var)2-10 RNAi* **(L)** co-expressing discs were dissected after 24 h of expression. *egr, apt RNAi* co-expressing discs **(O)** were even dissected after 48 h of *egr* expression. Note how knockdown of these negative regulators does not lead to increase in the JAK/STAT reporter activity in the JNK/AP-1 signaling domain. Discs were stained with DAPI to visualize nuclei. Scale bars: 50 µm

**Figure S4.3.**
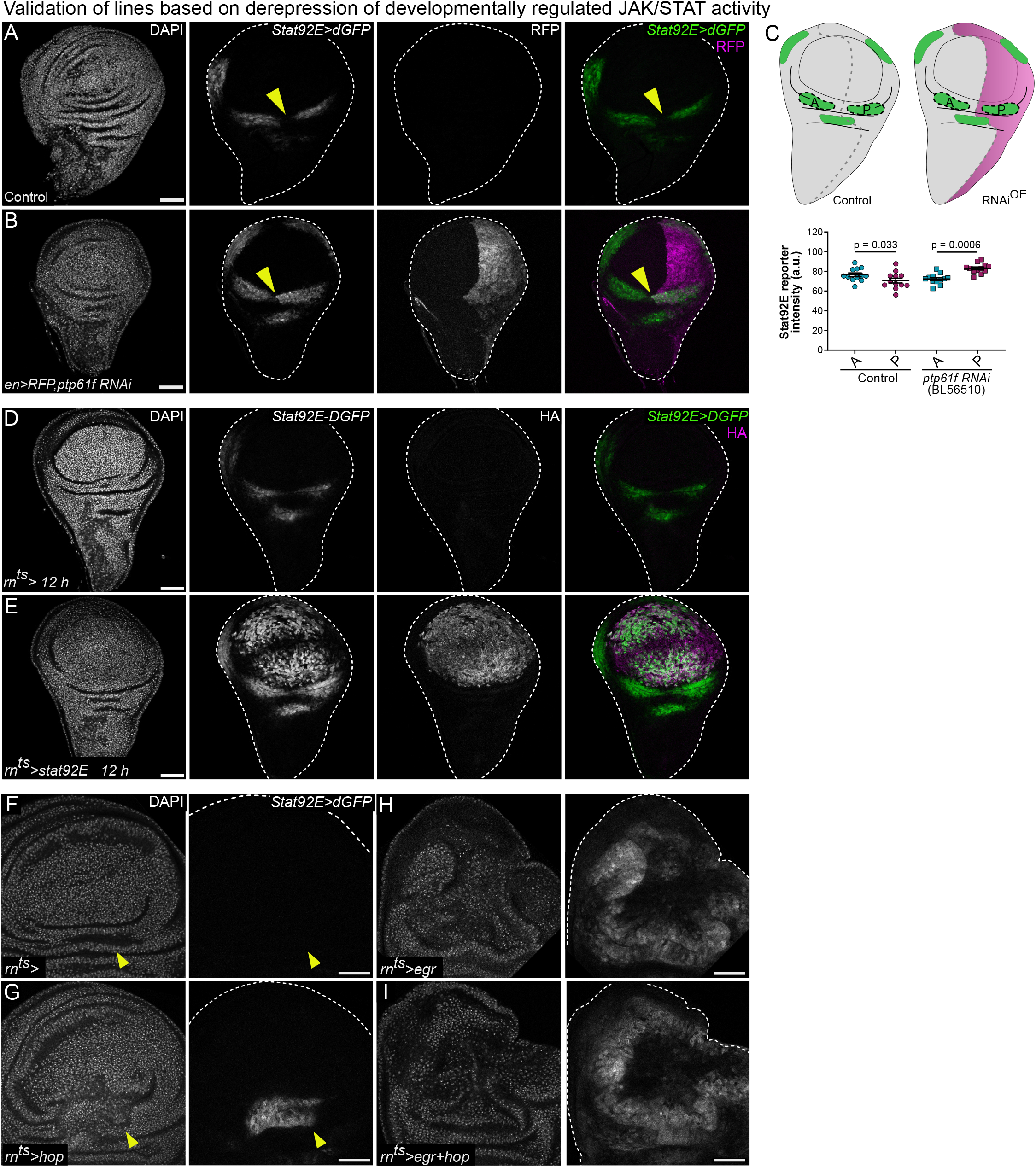
Testing the ability of *ptp61F-RNAi* and *UAS-Stat92E-HA* to elevate developmentally regulated JAK/STAT activity in the hinge or pouch. **(A-B)** Expression of *UAS-ptp61F-RNAi* (BL56510) and *UAS-RFP* (magenta) using the *en-GAL4* driver **(B)** leads to a distinct increase in developmentally patterned *Stat92E>dGFP* reporter (green) activity in the posterior wing hinge (yellow arrowheads) compared to stage-matched controls **(A)** dissected on D7 AED. **(C)** Quantification of changes to the development pattern of JAK/STAT activation, using fluorescence intensity measurements for the *Stat92E>dGFP* reporter. Black dashed lines in schematic show measured regions in the anterior (A) and posterior (P) wing hinge in control and *RNAi-*expressing (*en>RNAi*) discs. Graphs display mean ± SEM for *n=10,* controls and *n=12* RNAi-expressing discs. Paired t-tests were performed to test for statistical significance. **(D-E)** A Control **(D)** and a *UAS-STAT92E-3xHA* expressing disc, under the control of the *rn-GAL4* driver **(E)**, stained for HA (magenta). 12h of expression on D7 AED leads to strong upregulation of the *Stat92E>dGFP* reporter (green) in the wing pouch. **(F-I)** Compared to control discs **(E)**, overexpression of hop (UAS-hop.H) **(F)** leads to restricted, ectopic activation of the *Stat92E>dGF* reporter in the medial hinge fold (yellow arrowheads). However, compared to *egr*-expressing discs **(G)**, *egr,hop*-co-expressing discs **(H)** at R0 do not induce ectopic JAK/STAT activity in the central JNK/AP-1 signaling pouch domain. Discs were stained with DAPI to visualize nuclei. Scale bars: 50 µm

**Figure S5.**
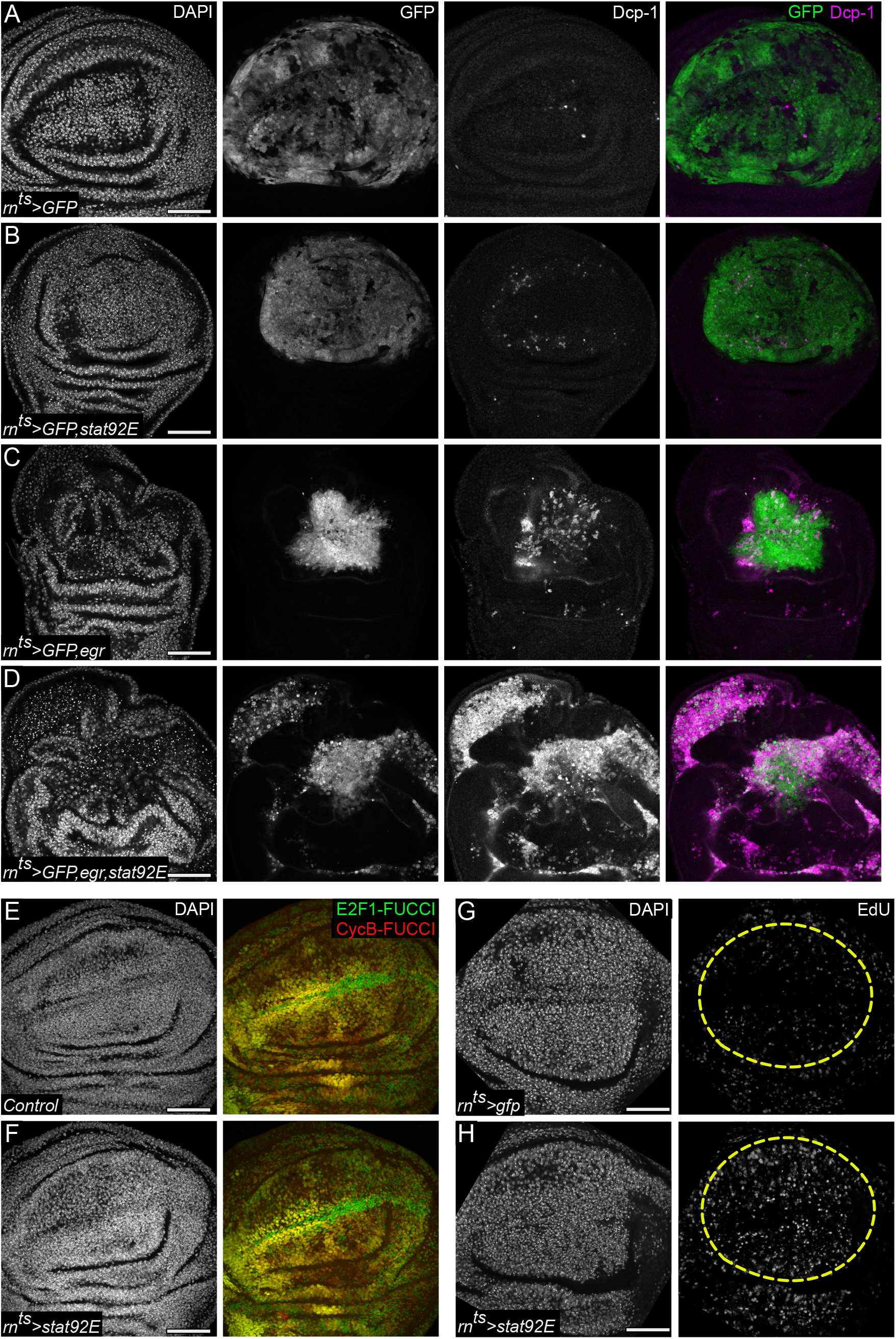
Cell-autonomous co-activation of JNK/AP-1 and JAK/STAT signaling gives rise to apoptosis. **(A-D)** A control wing disc **(A)**, *stat92E*-expressing **(B)**, *egr*-expressing **(C)** and *egr,stat92E*-co-expressing disc **(D)**, which also express UAS-GFP (green), all under the control of *rn-GAL4.* Discs were stained for cleaved Dcp-1 (magenta) to visualize apoptosis. Note how almost all apoptotic debris is labelled by GFP **(D)**, demonstrating that apoptotic cells truly originate from *egr,stat92E*-coexpressing cells. **(E-F)** A control **(E)** and *stat92E*-expressing wing disc **(F)** at R0, also expressing the FUCCI reporter. **(G-H)** A *gfp*-expressing control **(G)** and *stat92E*-expressing disc **(H)** at R0, assayed for S-phase activity by EdU incorporation. Elevated EdU is detected in the pouch domain ectopically expressing Stat92E. However, no changes were observed in the FUCCI cell-cycle profile. Maximum projections of multiple confocal sections are shown in **E-F** Discs were stained with DAPI to visualize nuclei. Scale bars: 50 µm

**Figure S6.**
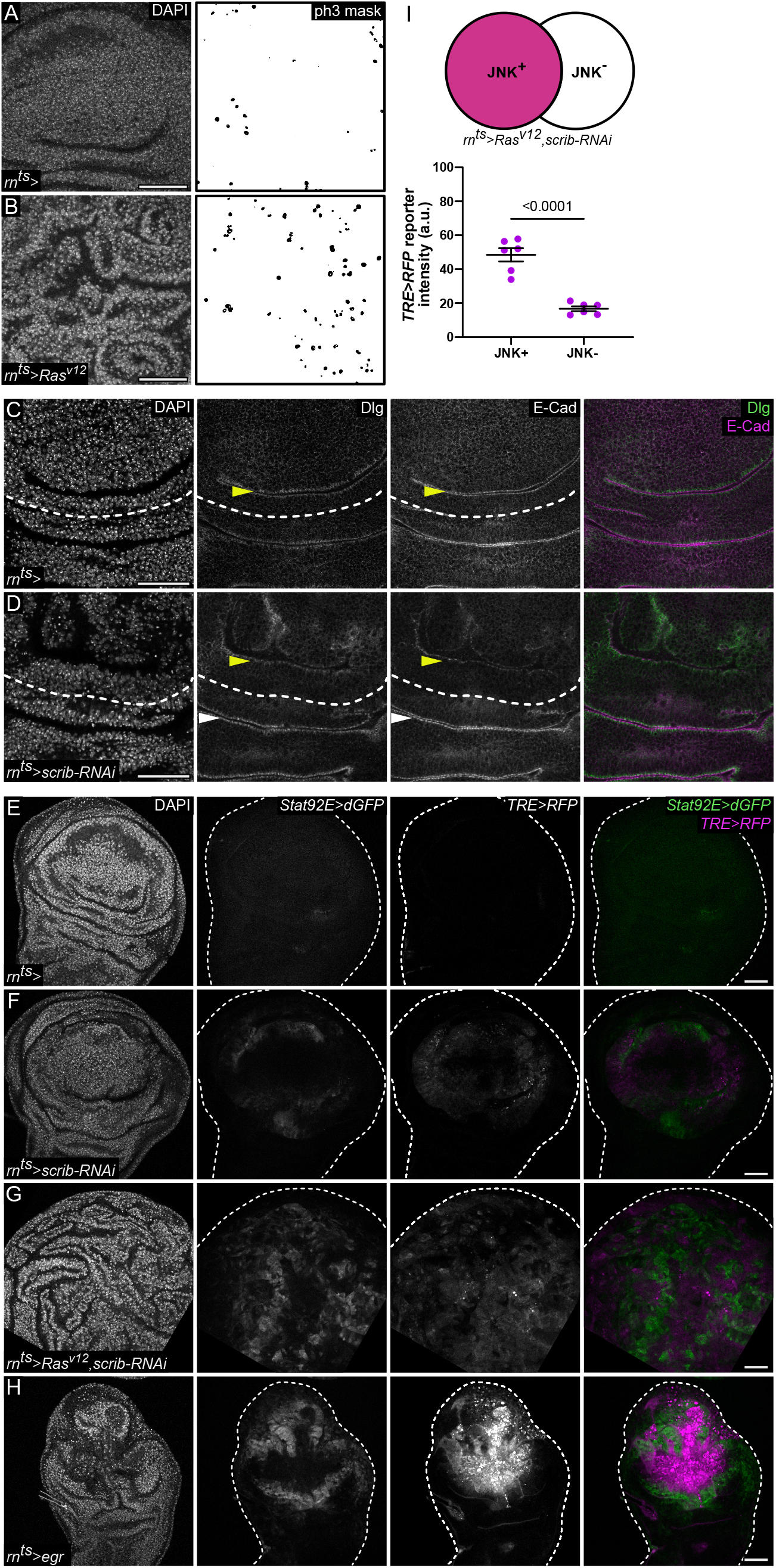
Tumors utilize lower but persistent levels of JNK/AP-1 activation to drive overgrowth. **(A-B)** A control wing disc **(A)** and wing disc expressing *Ras^v12^* **(B)** for 44 h. Discs were stained for phospho-Histone3 (ph3) to visualize cells in mitosis. Segmented masks of ph3 are shown. **(C-D)** A control wing disc **(C)** and wing discs expressing *scrib-RNAi* **(D)** for 44 h stained for E-cad (magenta) and Dlg (green) reveals loss of barrier integrity (yellow arrowheads in **D**) in the *scrib-RNAi*-expressing pouch domain when compared to its respective hinge domain (white arrowheads in **D**), as well as to the undamaged control pouch (yellow arrowhead in **C**). The pouch-hinge interface is marked with white dashed lines. **(E-H)** A control wing disc **(E)** and wing discs expressing *scrib-RNAi* **(F)**, *Ras^v12^*,*scrib-RNAi* **(G)** and *egr* **(H)** for 44h. Discs also express the JNK/AP-1 reporter *TRE>RFP* (magenta) and JAK/STAT reporter *Stat92E>dGFP* (green). *TRE>RFP* and *Stat92E>dGFP* reporter fluorescence intensities were adjusted to subsaturation in *egr*-expressing discs and all genotypes were imaged at comparable settings. **(I)** Schematic shows *TRE>RFP*-positive (JNK+, magenta) and *TRE>RFP*-negative (JNK-, white) regions selected for measurements. Graph represents *TRE>RFP* reporter fluorescence intensity measured within selected JNK+ and JNK-regions. Graphs represent mean ± SEM for *n=6, Ras^v12^,scrib-RNAi*-expressing discs. Paired t-tests was performed to test for statistical significance. Maximum projections of multiple confocal sections are shown in **A-B** Discs were stained with DAPI to visualize nuclei. Scale bars: 50 µm

**Table S1.**
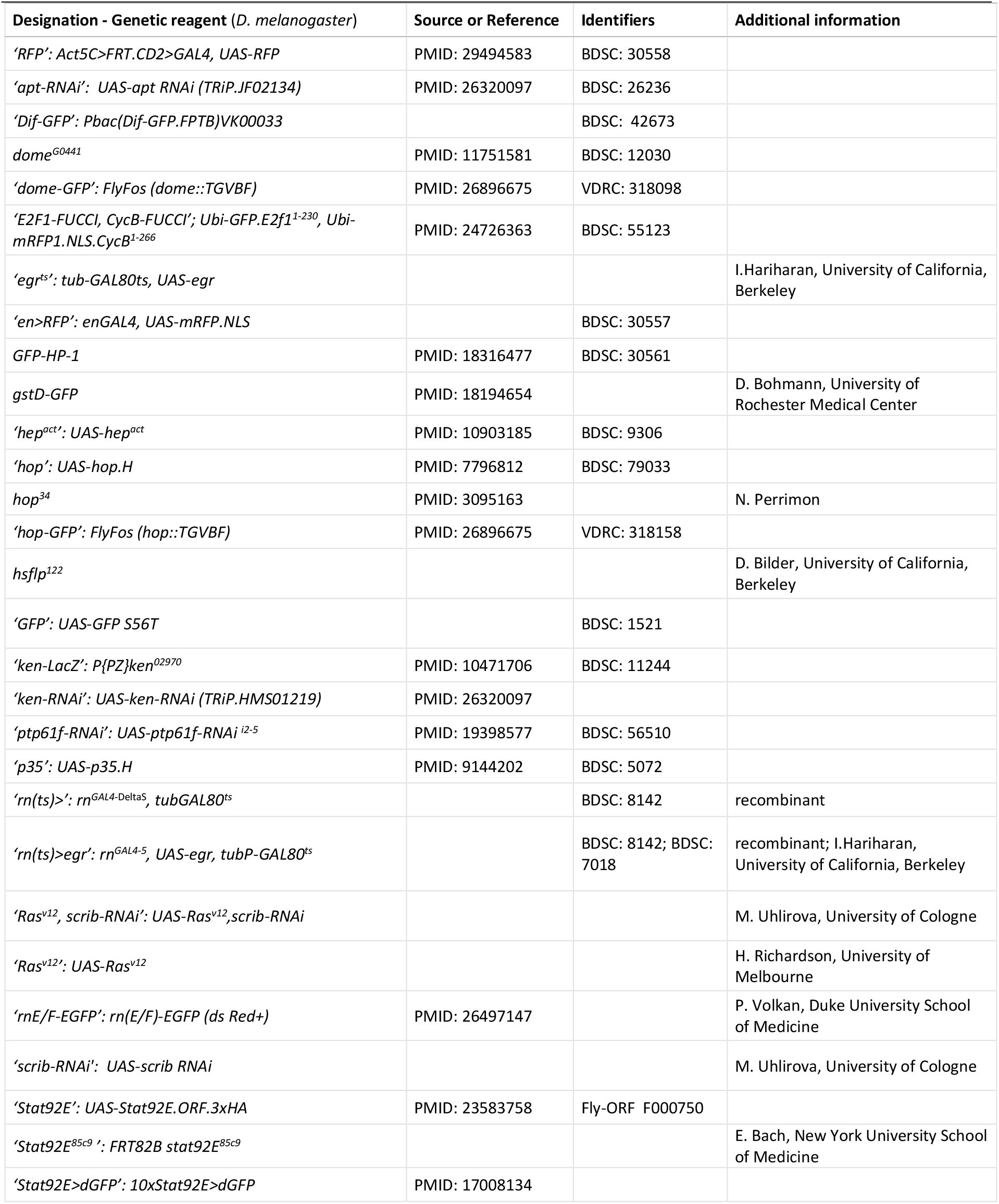
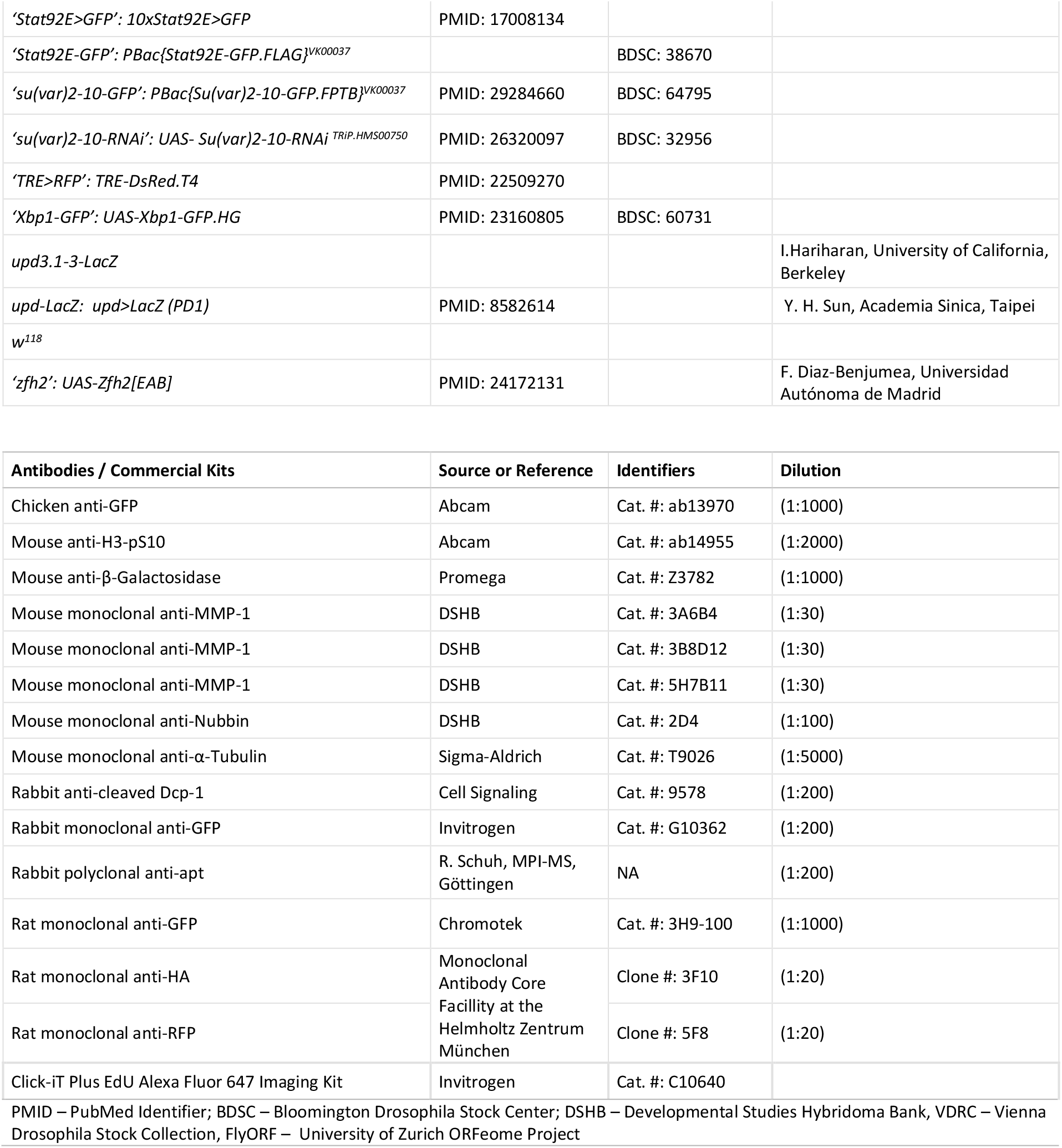
Key Resources Table

**Table S2.**
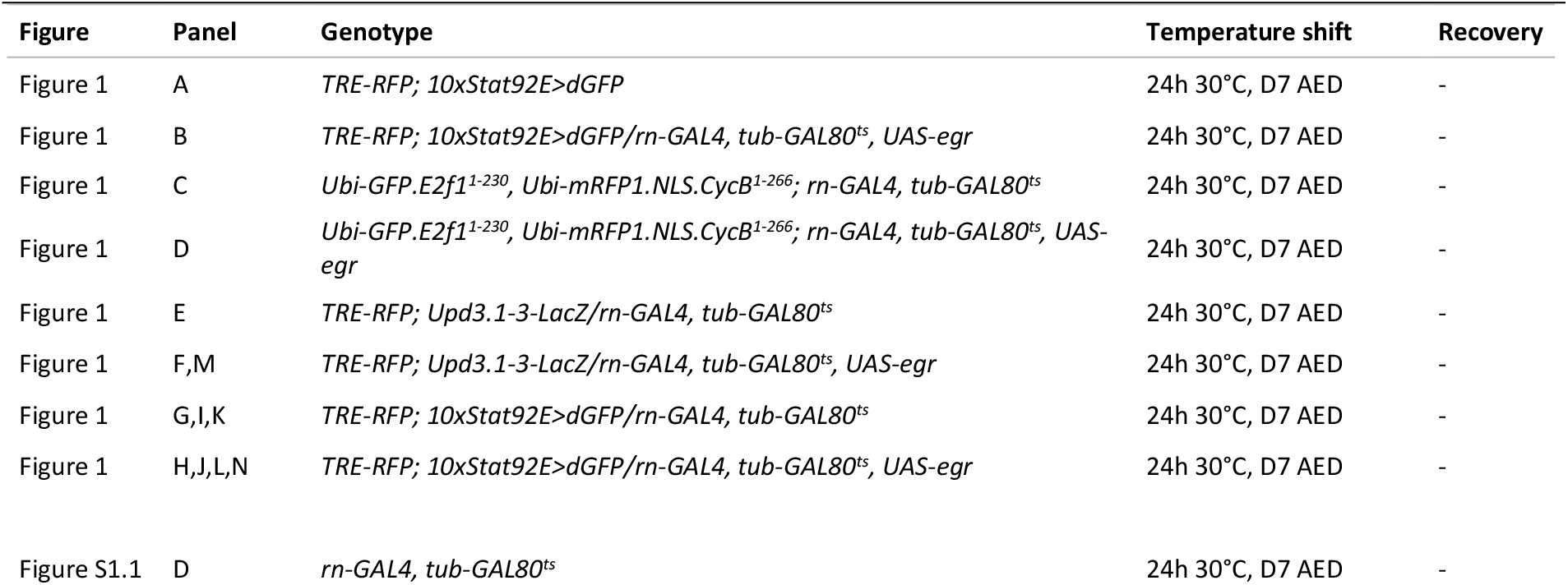
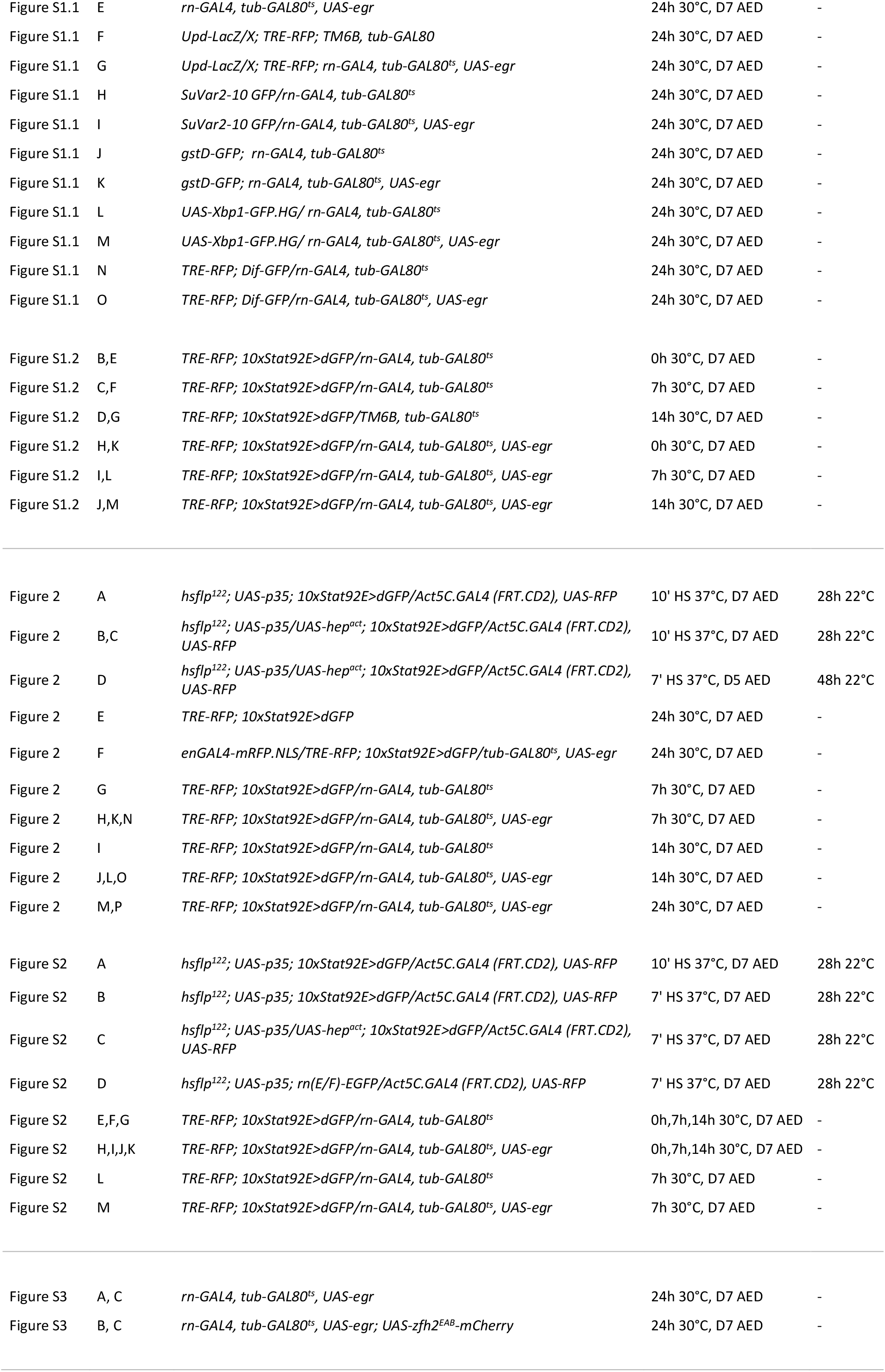
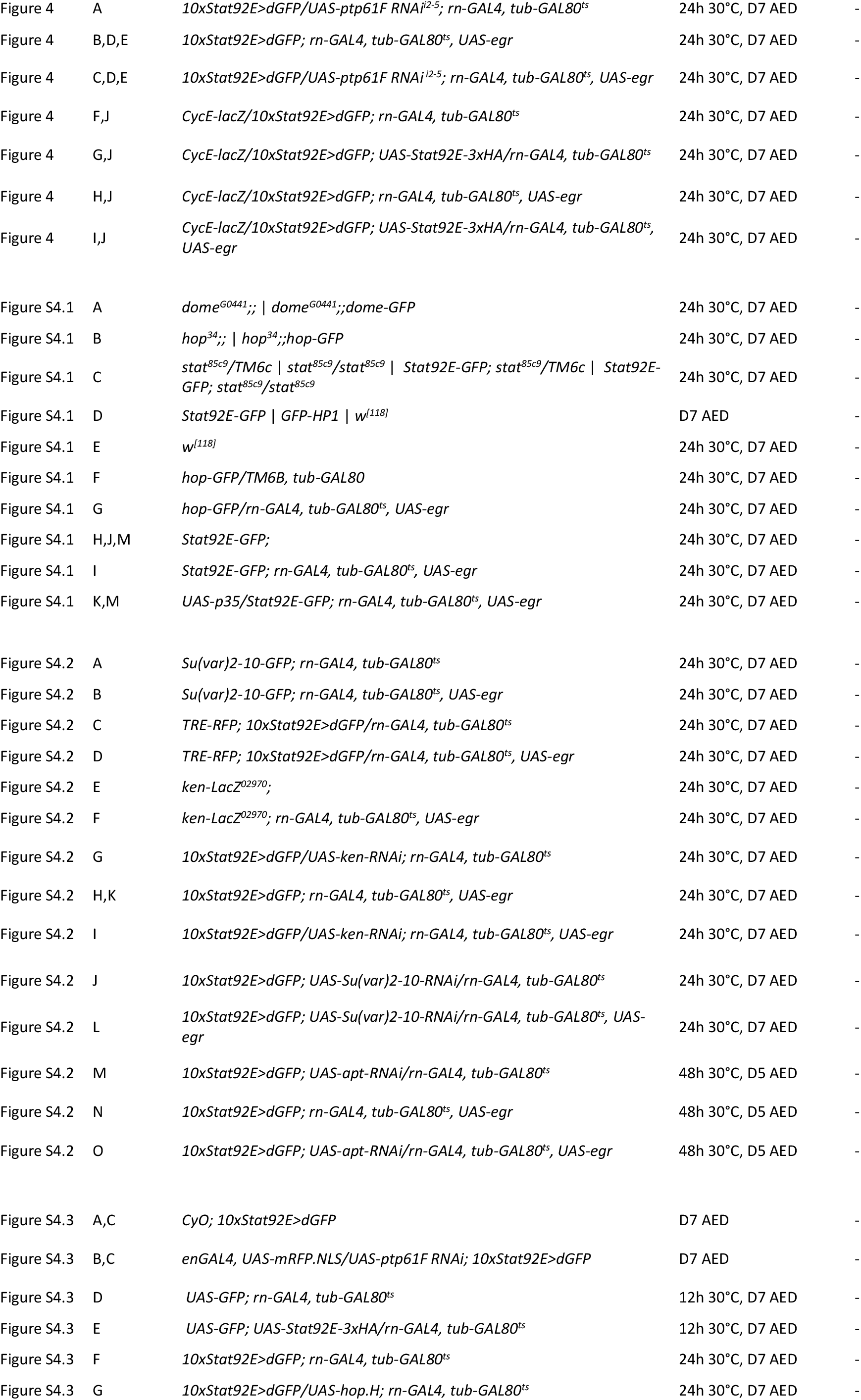
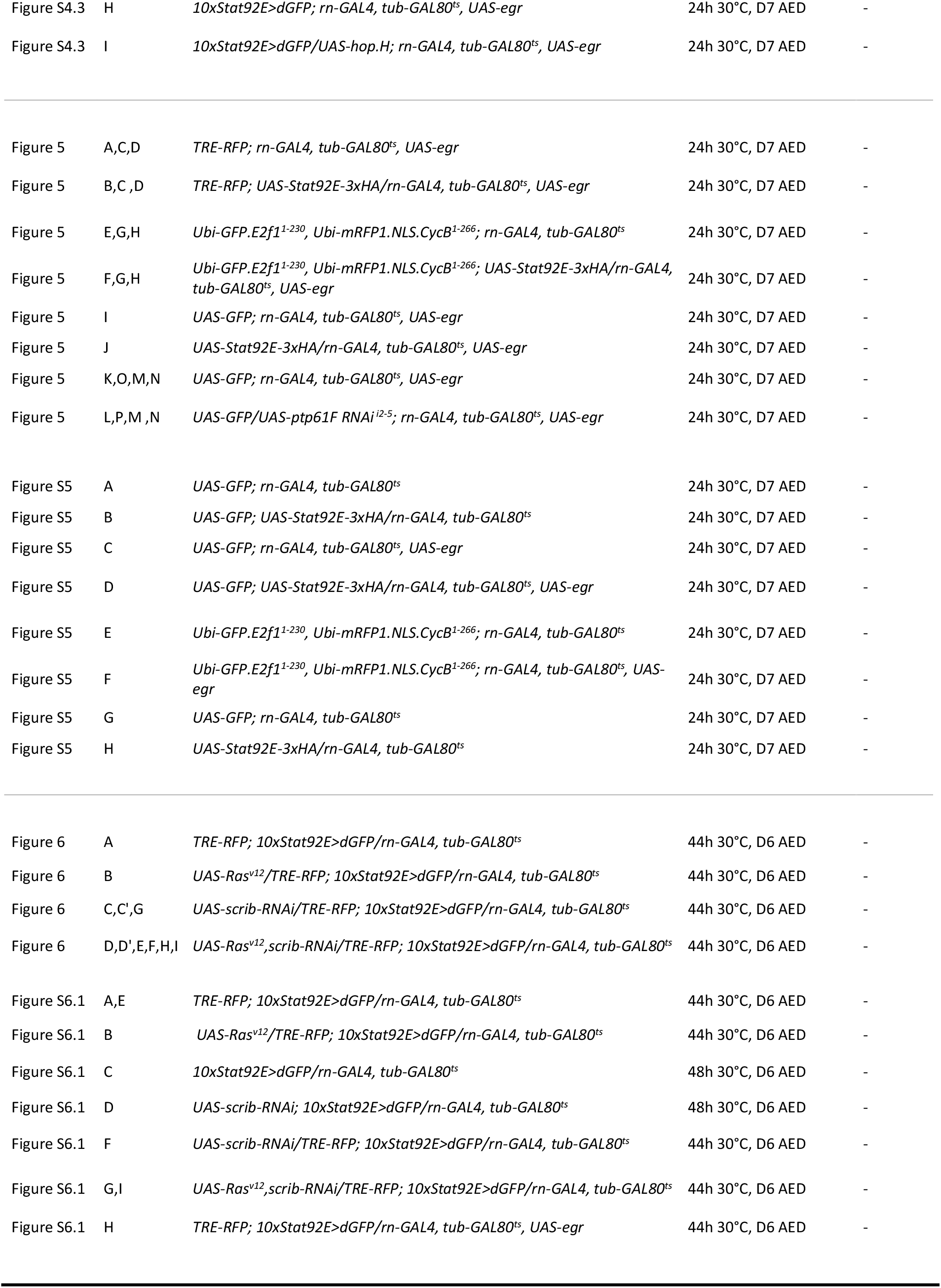
Detailed Genotypes

